# Comparing Neanderthal introgression maps reveals core agreement but substantial heterogeneity

**DOI:** 10.1101/2025.09.23.678138

**Authors:** Yaen Chen, Keila Velazquez-Arcelay, John A. Capra

**Affiliations:** Biological and Medical Informatics PhD Program, University of California, San Francisco, CA; Division of Hematology/Oncology, Department of Medicine, University of California, San Francisco, CA; Department of Bioengineering, University of California, San Francisco, CA; Bakar Computational Health Sciences Institute, University of California, San Francisco, CA

## Abstract

Statistical methods to identify Neanderthal ancestry in modern human genomes rest on varying assumptions and inputs. Nonetheless, most studies of introgression use only a single method to define Neanderthal ancestry. Due to a lack of “ground truth,” we have a limited understanding of the accuracy, comparative strengths and weaknesses, and the sensitivity of downstream conclusions for these methods. Here, we performed large-scale comparisons of genome-wide introgression maps from 12 representative Neanderthal introgression detection algorithms. These span methods that consider archaic and human reference genomes not from Africa (ArchaicSeeker2, CRF, DICAL-ADMIX), only archaic genomes (S*, Sprime, HMM, SARGE, ARGWeaver-D), only human reference genomes, including from Africa (IBDmix), or simulated data (ArchIE). Our results highlight a core set of regions predicted by nearly all methods, as well as substantial heterogeneity in commonly used Neanderthal introgression maps. Furthermore, we find that downstream analyses may result in different conclusions depending on the map used. Thus, we recommend careful consideration of map(s) chosen for an analysis and support the use of multiple maps to ensure robustness of conclusions. We make integrated prediction sets available, enabling further understanding of Neanderthal introgression’s legacy on modern humans.

## 1 Introduction

One of the closest extinct relatives of modern humans, the Neanderthals, lived across western and central Eurasia for hundreds of thousands of years before the ancestors of modern Eurasians migrated out of Africa^1^. Today, the Neanderthal legacy persists through artifacts, fossil remains, and Neanderthal genome sequences^2–4^. Analyses of Neanderthal and modern human genomes revealed multiple periods of interbreeding among the ancestors of modern humans, Neanderthals, and other ancient hominin groups, resulting in introgression—the exchange and integration of genetic information between Neanderthals and modern humans^5^. Most introgressed DNA observed in modern humans today originates from a substantial introgression event that occurred 40,000–50,000 years ago, while humans and Neanderthals co-existed^6,7^. The patterns of Neanderthal ancestry remaining in modern humans today after over ∼2,000 generations were shaped by natural selection, drift, and demographic processes. In modern humans not from Sub-Saharan Africa, 1-2% of each individual’s genome consists of Neanderthal ancestry^4,8^. Many methods are now available to identify Neanderthal introgressed DNA in modern humans, spanning a wide range of statistical approaches.

Neanderthal introgression detection methods can be broadly classified based on their genome inputs and statistical methods used. Probabilistic models, including hidden Markov models (HMMs)^9^ and Conditional Random Fields (CRF)^10,11^, rely on using a Neanderthal genome as an archaic reference and genomes from modern humans with limited Neanderthal introgression as outgroups. Commonly, Yoruba in Ibadan, Nigeria (YRI) from the 1000 Genomes Project (1KG)^12^ are used as the outgroup. HMM models that do not use an archaic reference have also been developed, while still using genomes from Africa as non-introgressed outgroups^12,13^.

Dynamic programming-based approaches, like the S* statistic^8^, identify sets of SNPs in high linkage disequilibrium (LD) that differ from an introgression-free outgroup and match expected lengths based on the timing of Neanderthal introgression. The S* approach assumes 1KG YRI genomes are introgression-free but does not rely on the archaic Neanderthal genome reference. The S* method was updated in Sprime^14^, which allows for lower levels of archaic introgression by removing the fixed-window approach and accounts for the frequency of the introgressed haplotypes, mutation rates, and recombination rates.

More recently, methods that do not rely on both archaic and modern human reference genomes have yielded insight into the history of Neanderthal introgression. IBDMix^15^ is a method that detects introgression using shared identity-by-descent (IBD) between modern human and Neanderthal genomes, computing a log-odds score indicating the likelihood of shared alleles due to IBD. Notably, IBDMix does not rely on an introgression-free modern human reference. As a result, this approach revealed small amounts of Neanderthal ancestry in genomes of individuals from Africa, likely due to modern humans carrying Neanderthal-introgressed loci migrating back to Africa from Eurasia. Other methods that are not explicitly trained on Neanderthal or reference non-introgressed genomes include ARGweaver-D^16^ and Speedy Ancestral Recombination Graph Estimator (SARGE)^17^, which identify introgression using ancestral recombination graphs (ARGs) to reconstruct the full genealogy of modern human genomes. Additionally, Archaic Introgression Explorer (ArchIE)^18^ uses a logistic regression model trained on simulated introgressed genotypes to infer introgressed regions, without the use of archaic or modern non-introgressed human reference genomes.

The output of these algorithms consists of predicted introgressed genomic regions and/or variants in modern individuals. We refer to these outputs collectively as introgression maps. These maps are commonly used to investigate evolution following admixture^8,11,13,15^ and the contribution of Neanderthal introgression to various modern human traits^19–24^. Most maps agree on the approximate proportions of Neanderthal ancestry present in modern populations. However, agreement between maps has not been comprehensively quantified. Some differences between introgression maps is expected due to true differences in introgression patterns between the individuals student, but the limited cross-method comparisons suggest more substantial differences^9,12,15,17,18,25^. For example, introgression deserts, or regions depleted of Neanderthal ancestry across individuals, are thought to have previously harbored introgressed alleles that were disadvantageous in modern humans and thus removed through negative selection^11,26–30^. Several studies have identified introgression deserts, but the criteria used vary, resulting in different sets of desert regions available for further analysis^8,11,13,15,25^. Introgressed loci have also been associated with various traits, including immune response, skin and hair pigmentation, skeletal structure, cognitive traits, and chronotype^19,31^. However, the choice of introgression map could influence downstream analysis of evolutionary processes that shaped Neanderthal introgression and how Neanderthal ancestry contributes to modern human traits.

Here, we compare representative Neanderthal introgression maps to understand variation in introgression predictions across modern human populations, facilitate further method development, and establish a resource to support the robustness of future results. Due to the lack of ground truth, we focus on identifying similarities and differences in available introgression maps at base-pair resolution and investigating the evolutionary and phenotypic properties of introgressed regions in each map. We anticipate that our conclusions will generalize to other introgression events from other groups or time periods, such as from Denisovans into modern humans, since similar methods are used to quantify introgression. Our results provide a resource for analyzing Neanderthal ancestry regions in the modern human genome, and a deeper understanding of Neanderthal introgression patterns will help reveal structural and functional differences between Neanderthals and modern humans that have yet to be fully uncovered.

## 2 Results

### 2.1 Introgressed loci differ in length and genomic coverage between maps

To quantify genomic patterns of existing introgression predictions, we integrated twelve published introgression maps across eleven algorithms (Table 1; Methods) and compared their genome-wide coverage.

**Table 1:**
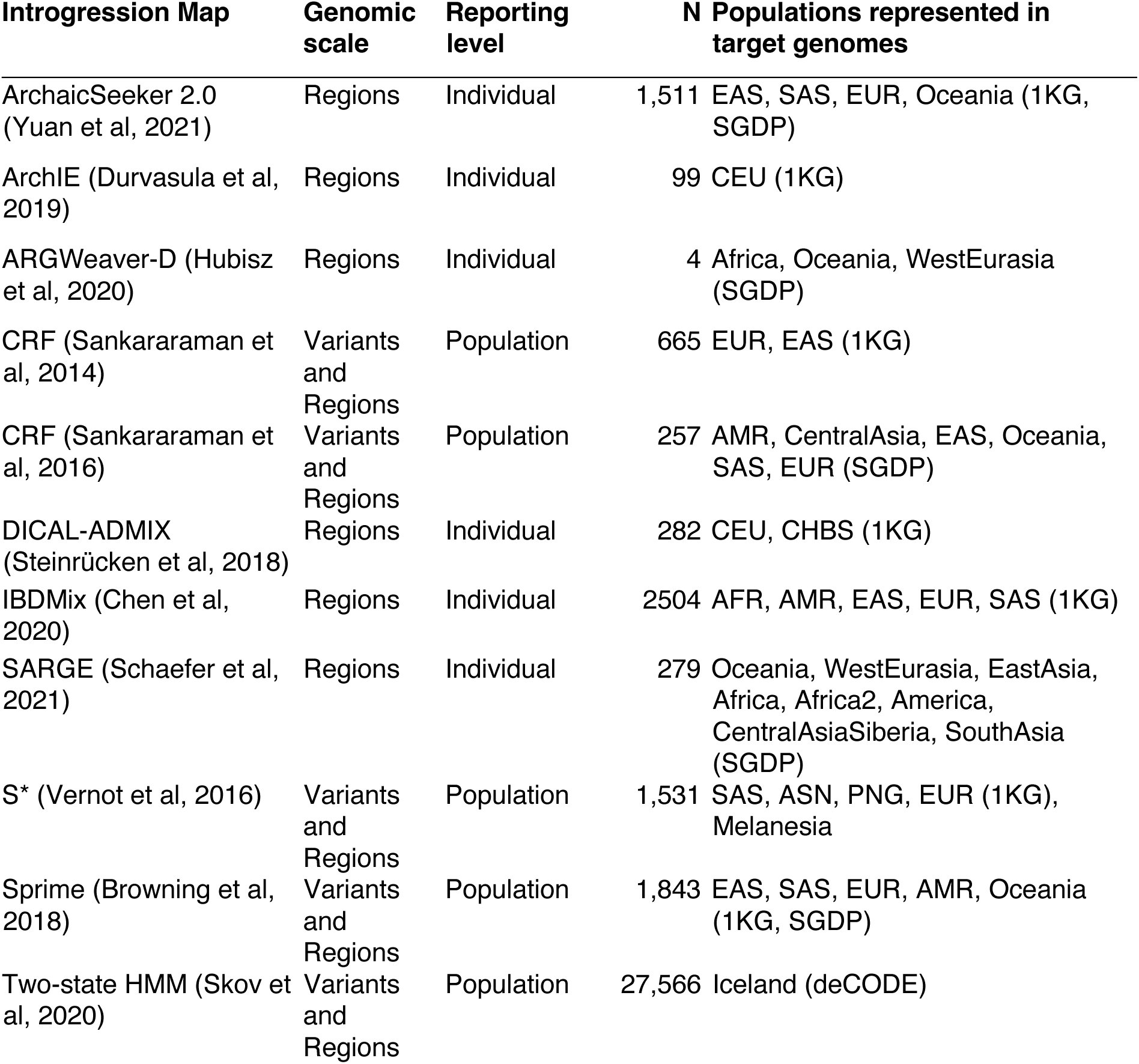
Properties of the introgression maps analyzed. . For each map, we describe whether Neanderthal introgression is reported at the region or variant level. Additionally, we indicate whether loci are provided at the individual-level or only as population-level introgression maps. Lastly, we describe the number of individuals included in individual-level analyses, as well as the number reported for each original study in population-level data. For Sankararaman et al 2016, we consider two sets of predictions: (1) using Simons Genome Diversity Panel (SGDP) African and Denisovan genomes as outgroups and (2) using only SGDP African genomes as an outgroup.

Overall, 2,242,601,207 bases, or ∼77% of the human genome (without assembly gaps), were called as introgressed in at least one map (Figure 2A). This is substantially greater than the proportion predicted by any single map; indeed, 17.1% of the introgressed base pairs were unique to a single map. The introgression maps also vary substantially in the lengths of the introgressed regions (Figure 2B). Given that these introgression maps are based on very different numbers of individuals and populations (Table 1), we caution that this first comparison is strongly influenced by the number and diversity of individuals considered by each method. For example, Sankararaman16 (1) which uses both SGDP African and Denisovan genomes as non-introgressed outgroups, identified the largest number of introgressed base pairs based on 257 SGDP non-African genomes. ARGweaver-D identified the smallest amount of introgression since it uses only four individuals, two of whom are from Africa, for introgression inference. The total number of base pairs detected as introgressed by a method begins to plateau after considering a few hundred individuals, though the dynamics vary across populations^33^. To mitigate the impact of the number of individuals considered in each introgression map, we focus in subsequent comparisons on the size-matched, highest-scoring predictions of each map or on predictions made in the same individuals. Due to the small number of individuals analyzed by ARGweaver-D, we excluded this method in map-wide comparisons.

**Figure 1:**
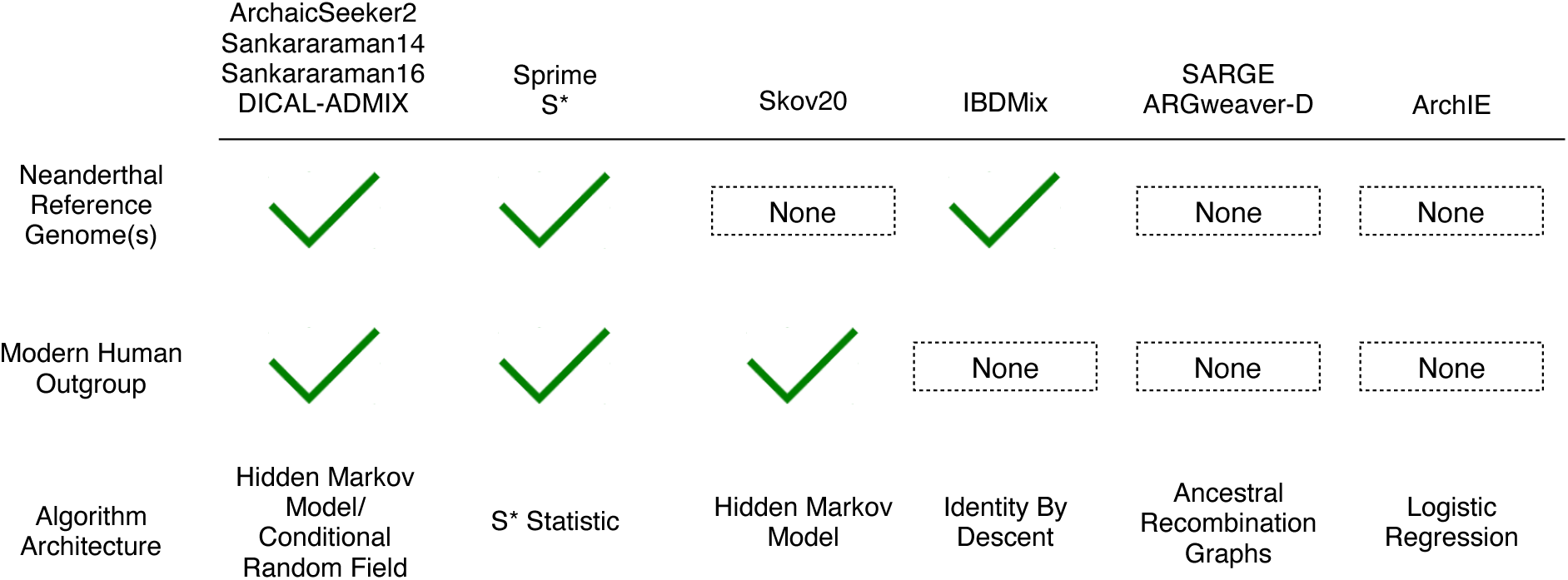
Classification of introgression detection methods analyzed based on their use of a Neanderthal reference genome, modern human outgroup, and underlying algorithm. Neanderthal reference genomes commonly include the high-coverage Altai^4^ and Vindija^3^ genomes. The modern human outgroups commonly consist of 1KG^32^ genomes from Yoruba. Algorithm architecture describes the underlying statistical approach for each method. See Table 1 for details.

**Figure 2:**
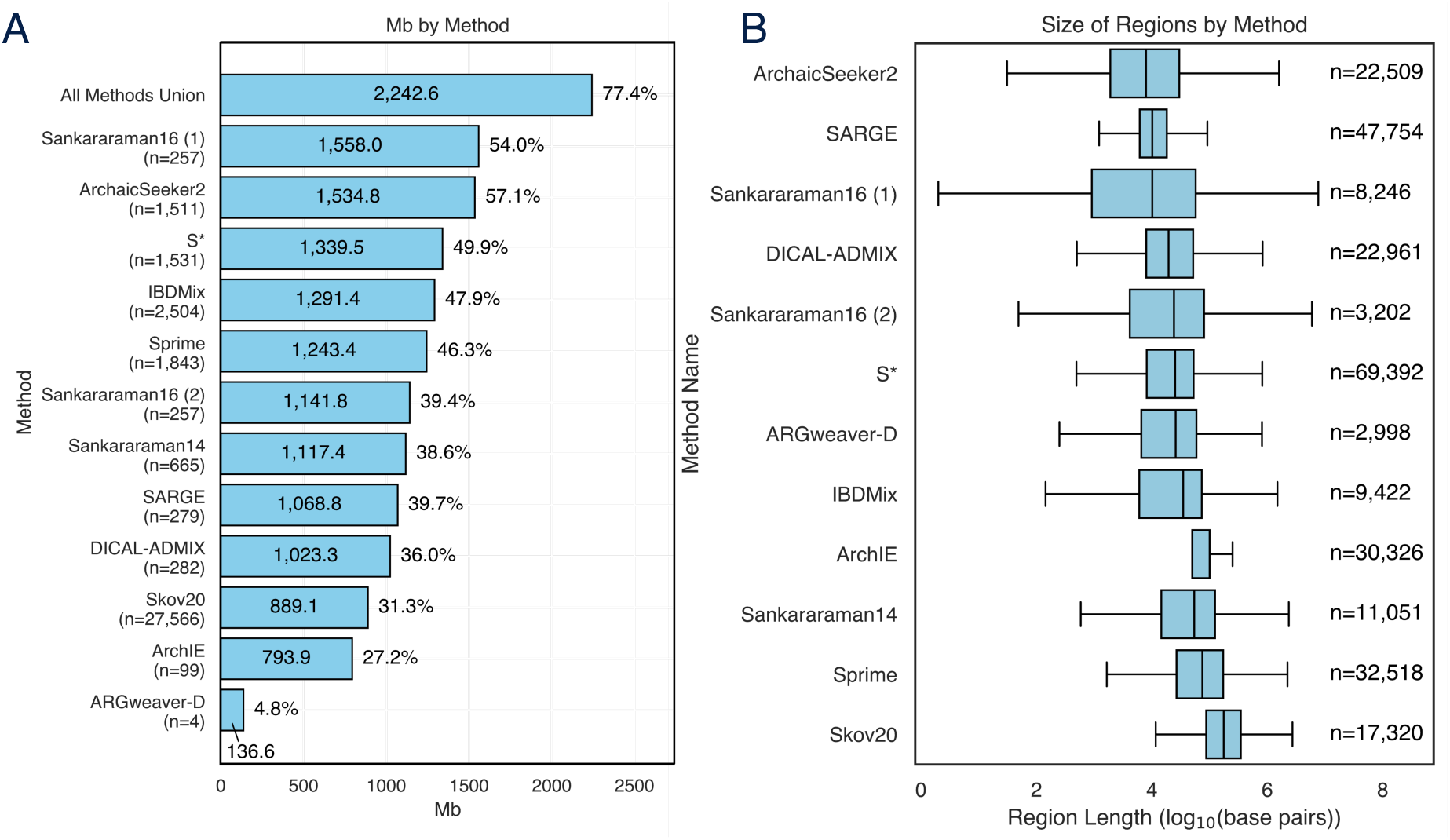
The genomic coverage and length distribution of each introgression map. (A) Amount of the genome (in millions of base pairs (Mb)) identified as having Neanderthal introgression in at least one individual by each map’s published predictions. Values reflect the union of introgressed loci across individuals and populations. Note that there are differences in the number and diversity of individuals analyzed that contribute to variation. Percentages outside each bar reflect genome coverage per map. (B) Boxplots of the distribution of introgressed region lengths (in base pairs) for each map. Ns to the right of each boxplot reflect the number of distinct, non-continuous regions for each map. Boxes reflect the interquartile range, and whiskers denote values outside 1.5 times the interquartile range.

### 2.2 Introgression maps have core agreement, but substantial discordance

Next, we quantified the number of maps that identified each genomic base pair as introgressed (Figure 3A,B). The most common scenarios are: 1) regions predicted as introgressed by all maps except ArchIE (197.65 Mb), 2) regions predicted only by ArchIE (151.54 Mb), and 3) regions predicted as introgressed by all maps (129.07 Mb). More broadly, regions were either predicted by many maps or only a few (Figure 3B). This suggests a core of regions identified by most introgression maps. We refer to loci supported by ten or more maps as highly supported regions in subsequent analyses (Figure 3B). Methods varied substantially in the amount of the genome they alone predicted as introgressed (Figure 3C).

**Figure 3:**
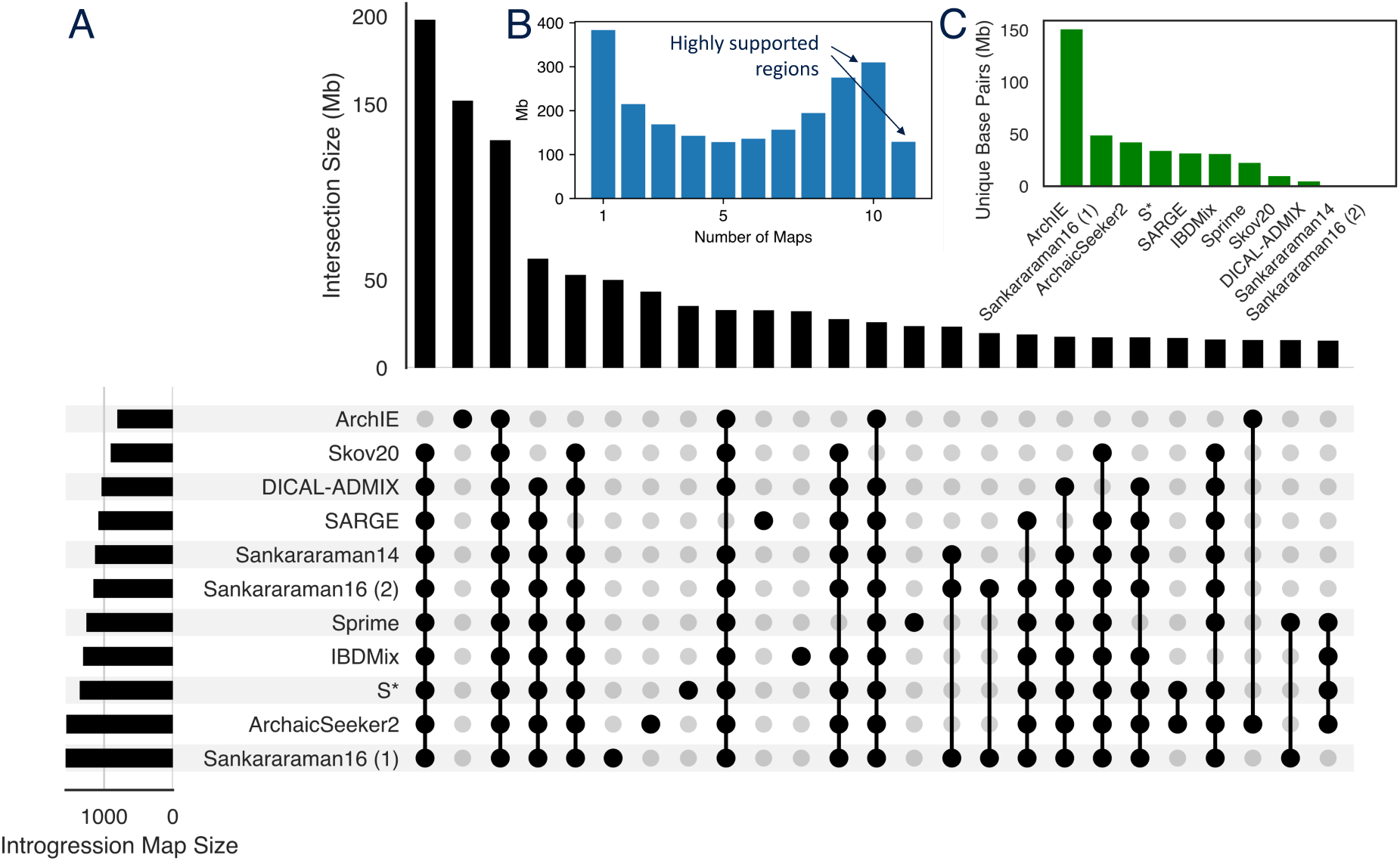
The overlap between introgression maps. (A) Upset plot of the amount of genomic overlap among regions identified by different combinations of introgression maps. The top 25 combinations are shown. (B) Histogram of the number of introgression maps supporting each bp predicted as introgressed. We refer to regions found in ten or more maps as “highly supported.” (C) The number of unique base pairs predicted as introgressed by each map.

To quantify the genome-wide agreement between introgression maps, we computed the Jaccard similarity—the size of the intersection divided by the size of the union—at the base pair level between pairs of maps. Additionally, we computed normalized Jaccard similarity, where the size of the smaller set is the denominator. These comparisons were dominated by the amount of the genome predicted as introgressed by a method (Supplementary Figure 19). Thus, to account for differences in the size of introgression maps, we reduced each introgression map to include only the highest-scoring regions matching the size of the smallest map (793.9 Mb from ArchIE) in subsequent analyses.

Overall, the maps showed substantial differences in their introgression predictions for autosomal regions (Figure 4). Raw Jaccard similarities ranged between 0.13 and 0.71 with an average of 0.31, highlighting that even the most similar maps have substantial differences in the regions they identify. The maps clustered into two main groups based on the Jaccard similarity: the CRF/HMM methods (Sankararaman14, Sankararaman16 (1) and (2), Skov20, DICAL-ADMIX) and the others (ArchaicSeeker2, Sprime, SARGE, IBDMix, S*). However, even after accounting for map size, ArchIE’s predictions were the most discordant with other methods.

**Figure 4:**
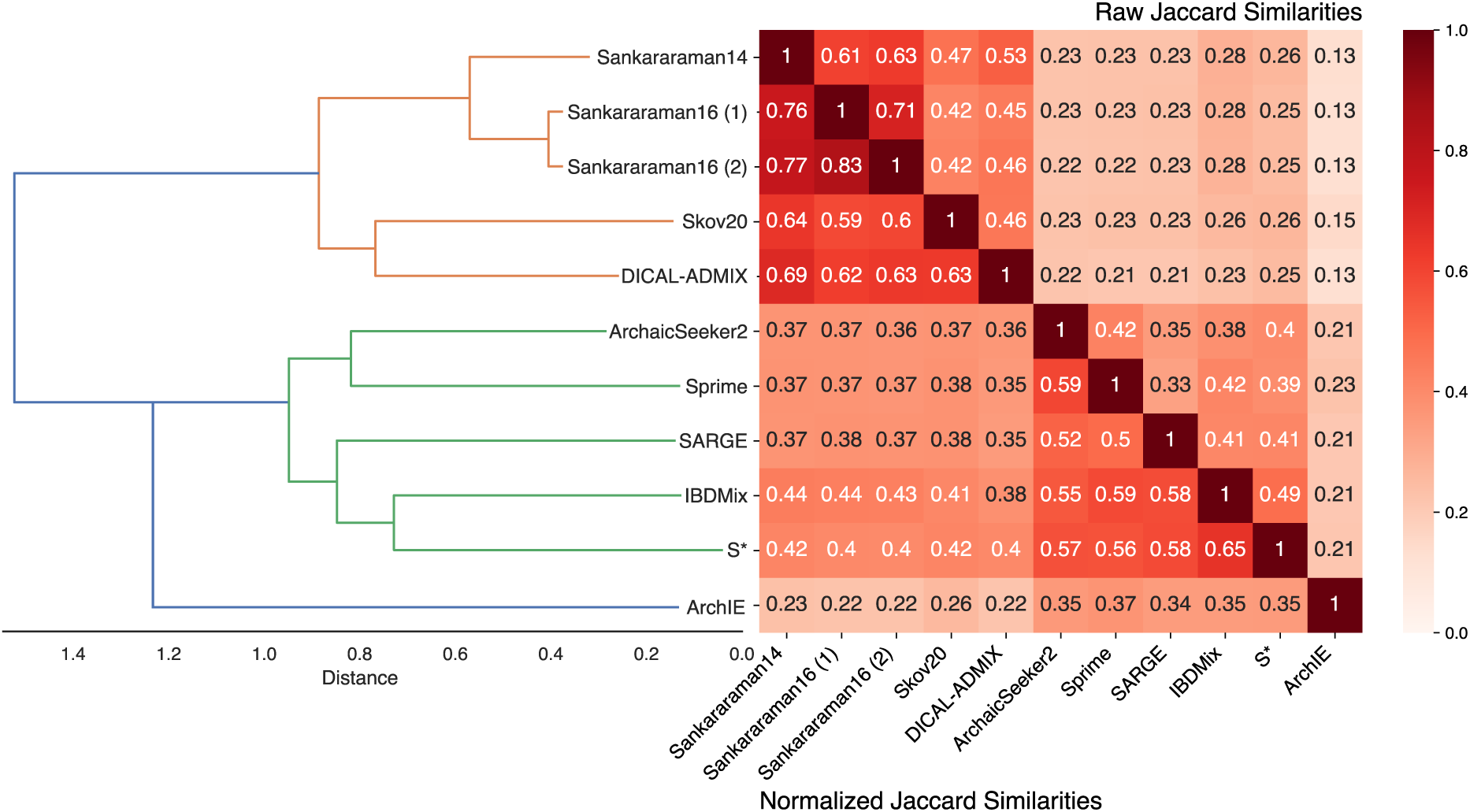
Jaccard similarities between top-scoring introgressed regions in autosomal chromosomes for each pair of maps. Jaccard similarities—the intersection divided by the union at the base pair level—are given above the diagonal. Normalized Jaccard similarities—the intersection divided by the size of the smaller set—are given below the diagonal. Introgression maps are ordered by hierarchical clustering based on their similarity, and distance values reflect the maximum distances between two sets, computed using farthest point clustering. To account for differences in the total amount of introgression predicted by different maps, this comparison is based on the top 793.9 Mb from each map. This amount was selected based on the amount of introgression predicted by the smallest map considered here (ArchIE). The comparison without this length constraint is dominated by the size of the map (Supplementary Figure 2).

Outside of autosomal regions, the X chromosome has been previously observed to be depleted in Neanderthal introgression due to potential hybrid incompatibilities^10^ and greater negative selection^30^. Only five out of 12 maps provide introgression predictions for the X chromosome: Skov20, Sankararaman16 (1) and (2), Sankararaman14, and DICAL-ADMIX. Introgression is depleted in the X chromosome compared to autosomes across methods: Each map has lower introgression coverage percentages in the X chromosome compared to autosomes. For introgressed regions in the X chromosome, agreement is more varied than autosomal-only comparisons after matching to the smallest set, Skov20 (Supplementary Figure 4). Introgression predictions in the X chromosome agree more when the comparison is not limited to top-scoring regions (Figure 5).

**Figure 5:**
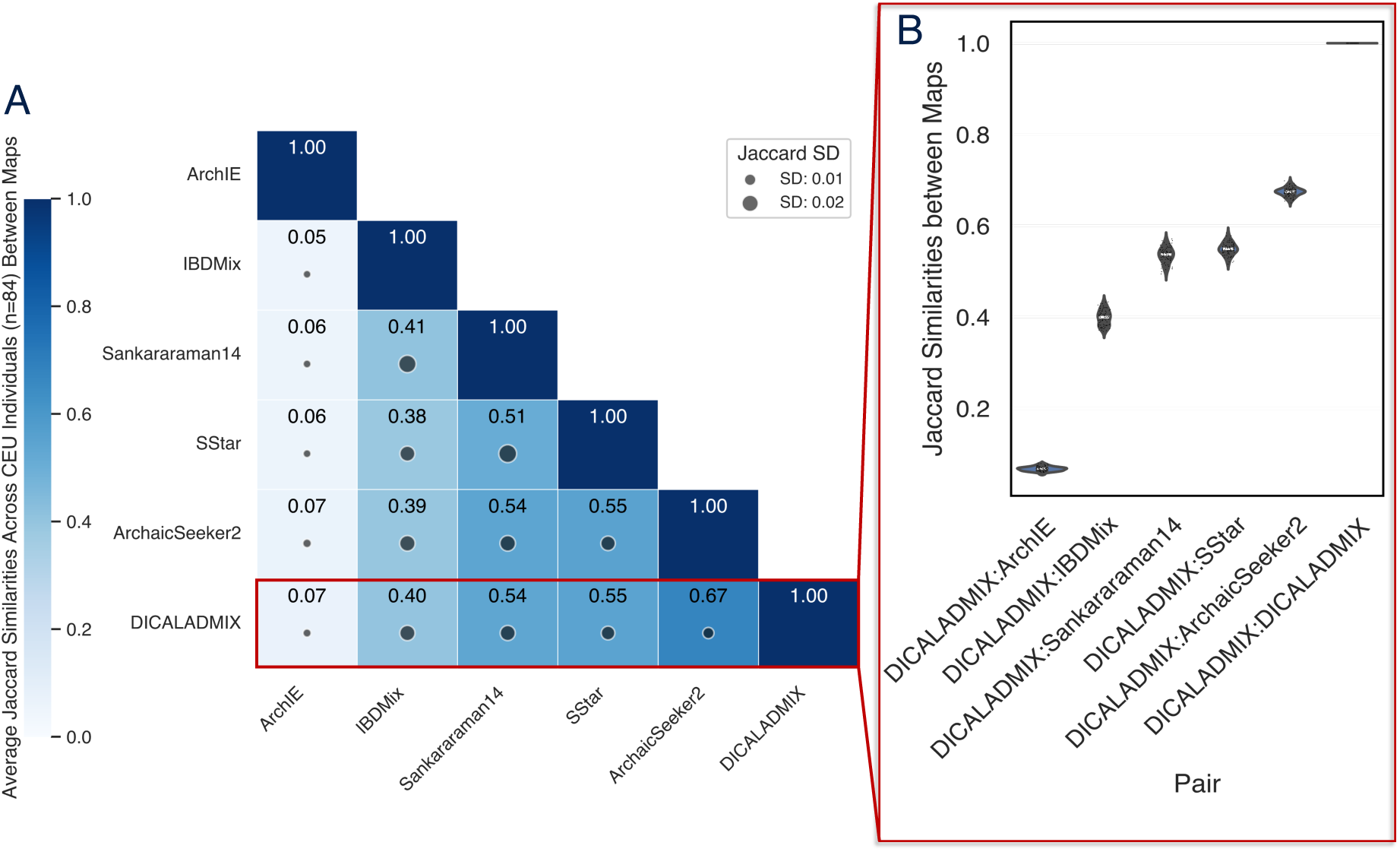
Comparison of introgression maps at the individual level. (A) Average Jaccard similarities between individual-level introgression maps for 84 individuals from Utah with Northern European ancestry (CEU 1000 Genomes population). Each cell gives the average Jaccard similarity for a pair of individual-level introgression maps for each individual. The size of the dot indicates the standard deviation of the 84 Jaccard similarities for each cell. (B) Violin plot of pairwise Jaccard similarities distributions over the 84 shared CEU individuals for each introgression method pair for the in the red highlighted row from (A).

In addition to region-level introgression maps, six methods call specific Neanderthal introgressed variants: Sankararaman16 (1) and (2), Sankararaman14, Skov20, Sprime, and S*. At this variant level, the Sankararaman variant maps all cluster together with Jaccardsimilarities ranging from 0.3-0.66, separate from Skov20, Sprime, and S*, which form a separate cluster with Jaccardsimilarities ranging from 0.12-0.37. In this group, S* and Sprime have the highest agreement (Jaccard=0.37) (Supplementary Figure 8). Four methods identify variants in the X chromosome: Skov20, Sankararaman (1) and (2), and Sankararaman14. X chromosome introgressed variants have the highest agreement among Sankararaman methods (Jaccard=0.61) (Supplementary Figure 10).

### 2.3 Individual-level comparisons highlight discordance between introgression maps

Next, we investigated how introgression maps from different methods vary in their predictions at the individual level. We focus on autosomal introgressed regions predicted on several sets of individuals. The first group consists of 84 individuals from Utah with Northern European ancestry (CEU individuals) from the 1KG, with introgression predictions from six methods (Figure 5).

These comparisons reveal that different introgression detection methods often yield substantially different predictions when applied to the same individual’s genome, with Jaccard similarities ranging from 0.05 to 0.67 (Figure 5). ArchIE’s maps are also the most discordant at the individual level, with an average Jaccard with other maps lower than 0.1. IBDMix was the next most discordant map, but it had much higher agreement with other maps with Jaccard statistics ranging from 0.39-0.41. Even though DICALADMIX and ArchaicSeeker2 were in different clusters in the genome-wide comparison, their individual-level introgressed region maps agreed more than any other pair of methods. We did not include maps that did not include predictions on these individuals.

For other groups of individuals with predictions from multiple methods (One SGDP Papuan individual, 228 SGDP individuals), we also observed similar patterns of moderate agreement between methods (Supplementary Figures 12, 13.

We also analyzed agreement between maps of introgressed regions in African genomes, which included IBDMix, ARGWeaver-D, and SARGE (Supplementary Figures 14, 15. The similarities between all pairs of maps were extremely low (average Jaccard similarity of 0.05, SD=0.0043). Given the small amounts of Neanderthal introgression and variation in numbers of African individuals between maps, the normalized Jaccard across these individuals’ maps is higher (Jaccards 0.33-0.44). Introgression prediction in genomes from Africa remains a challenge due to their common use as an outgroup and the small levels of Neanderthal ancestry present. Our limited comparisons reveal discordance between maps in introgressed regions from African individuals.

### 2.4 Introgressed regions in all maps have less evidence of background selection compared to regions without introgression

Analyses of the first Neanderthal introgression maps found that regions with high levels of Neanderthal ancestry have less evidence of background selection from linked functional elements than regions with low levels of Neanderthal ancestry^10^. As a first step in exploring the robustness of conclusions about patterns of Neanderthal ancestry, we tested this result across the introgression maps considered here. We computed the average B statistic, which quantifies the expected fraction of neutral diversity that is present at a site, across regions with introgression versus those without introgression for each map. Lower B values indicate stronger background selection^34^.

Each introgression map had significantly less evidence of background selection (higher average B values) in regions with predicted introgression compared to regions without inferred introgression (Figure 6). Sprime and ArchIE had the largest differences (ratio of averaged introgressed vs. non-introgressed B values: 1.069 and 1.061, respectively), while SARGE and ARGweaver had the smallest differences (ratio of averaged introgressed vs. non-introgressed B values: 1.019 and 1.004, respectively). Thus, all introgression maps support this conclusion and suggest selection against Neanderthal alleles in regions with many constrained functional elements.

**Figure 6:**
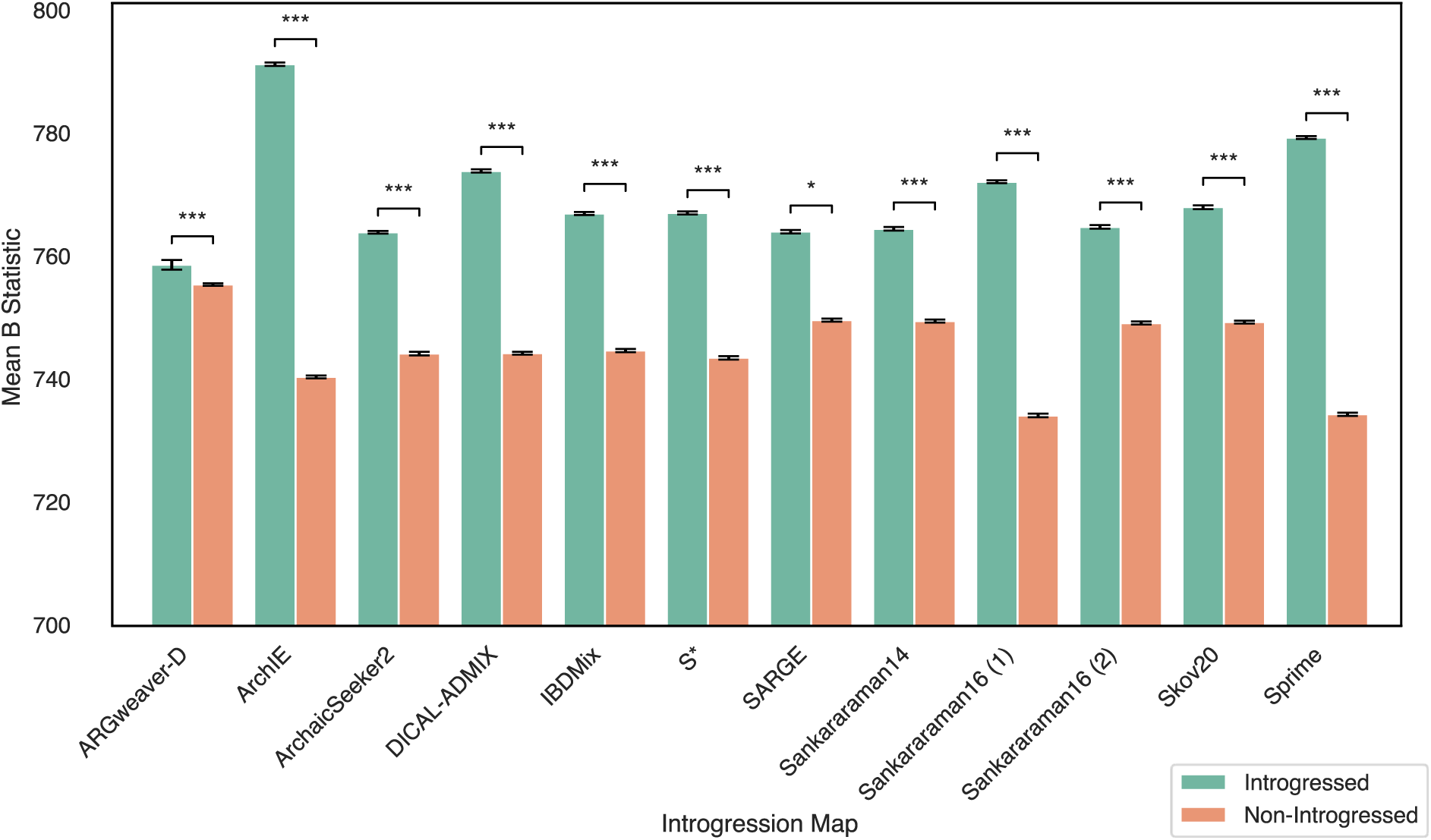
Neanderthal regions from each introgression map have less evidence of background selection. Average background selection scores as quantified by the B statistic^34^ for introgressed and non-introgressed regions from each map. B values were quantized in 500 bp bins following Telis et al.^27^ Higher B values indicate less background selection. Error bars reflect 95% confidence intervals based on 1,000 bootstraps.Asterisks denote P values: *P < 0.05, **P < 0.01, and ***P < 0.001 for Mann-Whitney U tests comparing B statistic values between introgressed and non-introgressed windows.

### 2.5 Phenotype annotations enriched in different introgression maps are discordant

Gene set annotation enrichment analysis is commonly performed on genes in regions with or without evidence of introgression. These analyses have yielded hypotheses about properties of the genome that retain Neanderthal ancestry versus regions that do not. For example, genes in regions with introgressed Neanderthal DNA have been associated with immune function, skin and hair, metabolism, and ultraviolet radiation sensitivity^8,15,26^. Some of these enrichments may reflect benefits of Neanderthal ancestry in regions encoding traits relevant to adaptation to Eurasian environments and pathogens^35^. However, further analyses are required to interpret the causes of phenotypic enrichment or depletion.

To test how robust annotation enrichment results are to different introgression maps, we performed gene set enrichment analyses on each introgression map and compared the results. We used rGREAT, a tool that performs gene annotation enrichment analysis by assigning input genomic regions to genes based on their proximity. rGREAT accounts for different probabilities of regions being assigned to a gene based on their length and distribution across the genome. For each introgression map, we computed enrichment compared to the whole genome background for gene functions from the Human Phenotype Ontology (HPO) (Supplementary File 1, 2).

The number of significantly enriched phenotypes for each introgression map varies substantially—from 0 to 734 (Figure 7). Overall, we discovered 1,676 significant enrichments (representing 1,354 unique phenotypes) between a phenotype and an introgression map. However, 1,090 of the enriched phenotypes are observed for only one introgression map. Several maps—ArchIE, IBDMix, Sankararaman16 (2), Skov20, and Sprime—do not have any HPO phenotype enrichments.

**Figure 7:**
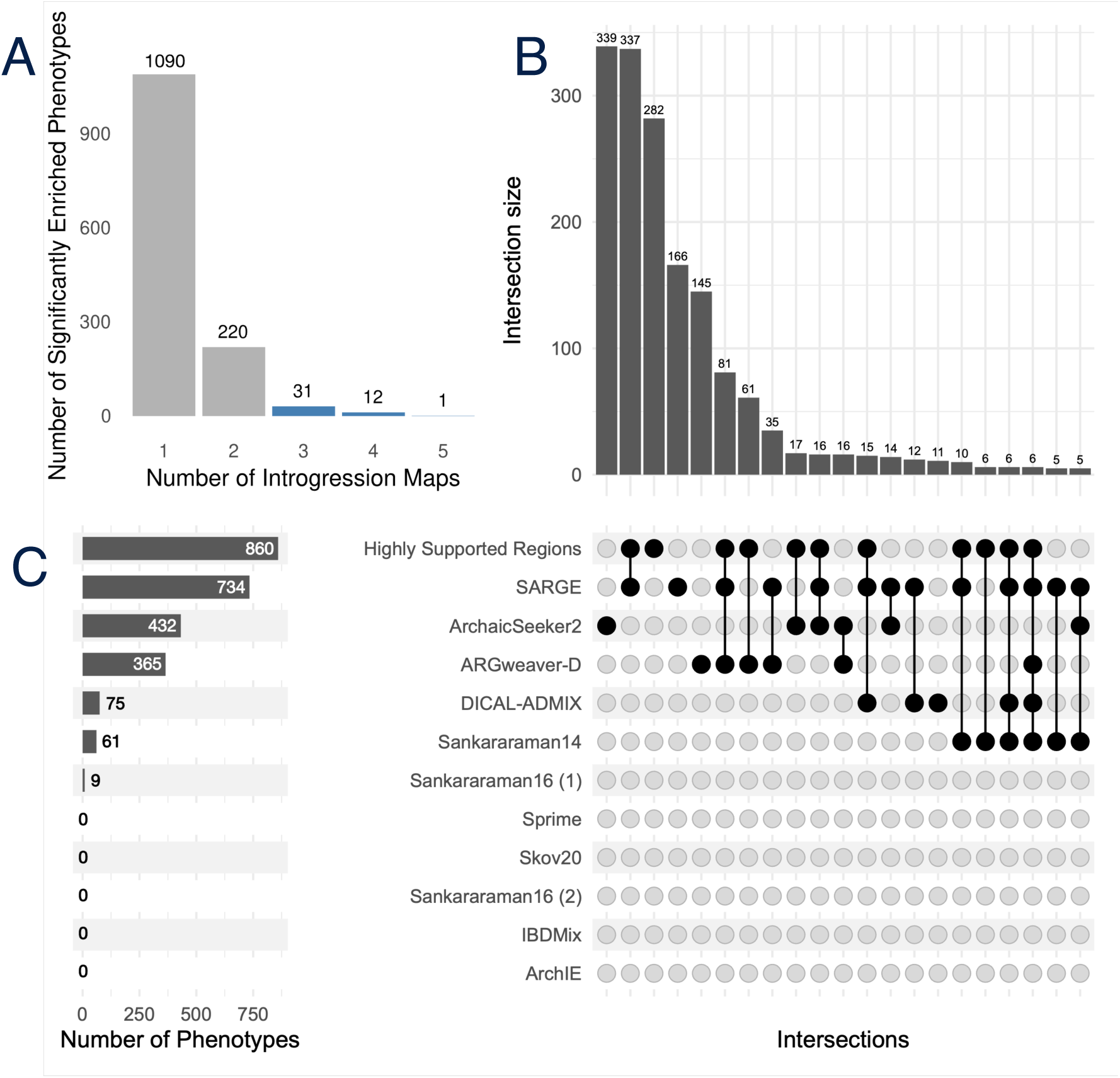
Different phenotypes are enriched in different introgression maps. (A) Number of significant Human Phenotype Ontology (HPO) phenotypes enriched in different numbers of introgression maps. Bars in blue reflect phenotypes supported by 3+ maps, which are represented in Figure 8. Gene set annotation enrichment for each introgression map was performed using rGREAT with gene annotations from the HPO. The significantly associated phenotypes for each method and highly supported regions are given in Supplementary File rGREAT_sig_phenos_include_highsupportloci.tsv. (B) Upset plot of HPO phenotypes supported by varying combinations of introgression maps. (C) Number of HPO phenotypes associated with each introgression map from (B), along with highly supported regions present in 10+ maps.

We identified 44 phenotypes enriched in at least three distinct maps (Figure 7). Many of these shared phenotypes influence a few specific bodily systems, including the eyes, skull, limbs, muscle, and reproductive tract. Patterns of similarity among the enrichments between maps differed from their overall genomic similarity (Supplementary Figure 17).

To explore whether the highly supported introgressed loci had similar enrichments to those found across multiple methods, we performed rGREAT phenotype enrichment analysis on sets of highly supported regions (Supplementary Table 1). These regions have a higher number of phenotypes associated (up to 965) compared to each introgression map individually. Moreover, 26 of the 44 phenotypes associated with 3+ maps are present in the set of phenotypes associated with highly supported regions.

To explore the robustness of these observations to other phenotype enrichment methods, we also carried out phenotype enrichment analyses using a stricter criterion for mapping introgressed regions to genes, requiring that the introgressed region overlaps a gene’s exon. Based on these gene sets, we used Annotatr and Enrichr to perform gene-set enrichment analyses on phenotypes from the GWAS Catalog 2023 and HPO. This stricter mapping resulted in fewer phenotypic associations, but the heterogeneity of the associations and a greater number of enrichments for the highly supported introgressed regions remained (Supplementary Table 2, Supplementary Figure 18).

### 2.6 Desert regions vary in levels of introgression across maps

Long genomic regions with very low levels of Neanderthal introgression have been identified in several previous introgression maps (Figure 8A). However, the criteria used to call deserts varied substantially (Supplementary Table 3), complicating comparisons and interpretation. To evaluate the support for five sets of deserts called in previous studies provided across introgression maps, we intersected all maps with desert regions to quantify levels of introgression across maps in the deserts (Figure 8B).

**Figure 8:**
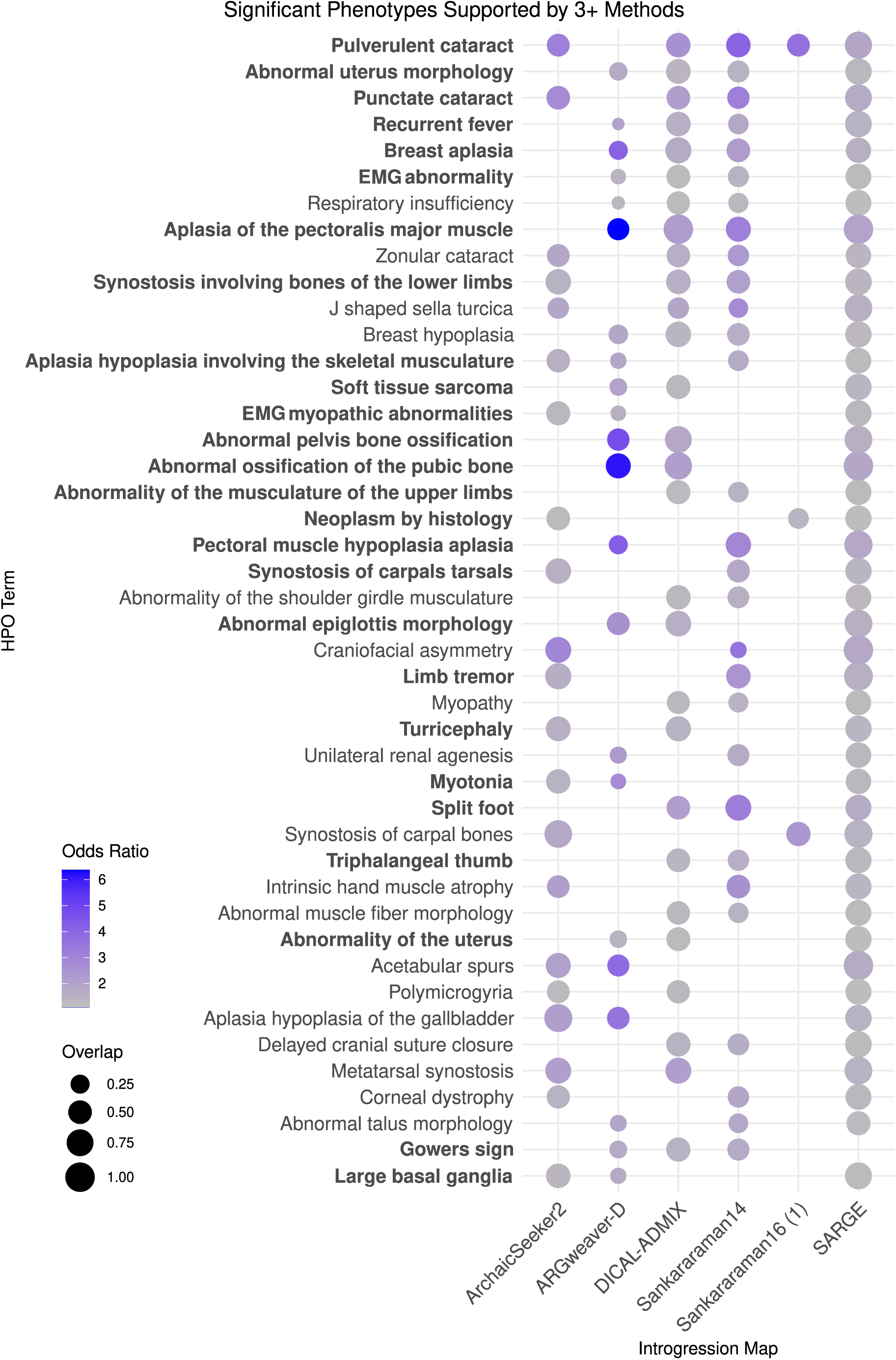
HPO phenotypes significantly supported by at least 3 methods’ introgression maps. Phenotypes are ordered by the number of introgression maps supporting enrichment, then by phenotypes with the lowest p-value across tests. 28 phenotypes highlighted in bold are also enriched in highly supported introgressed regions (Supplementary Table 1).

**Figure 8:**
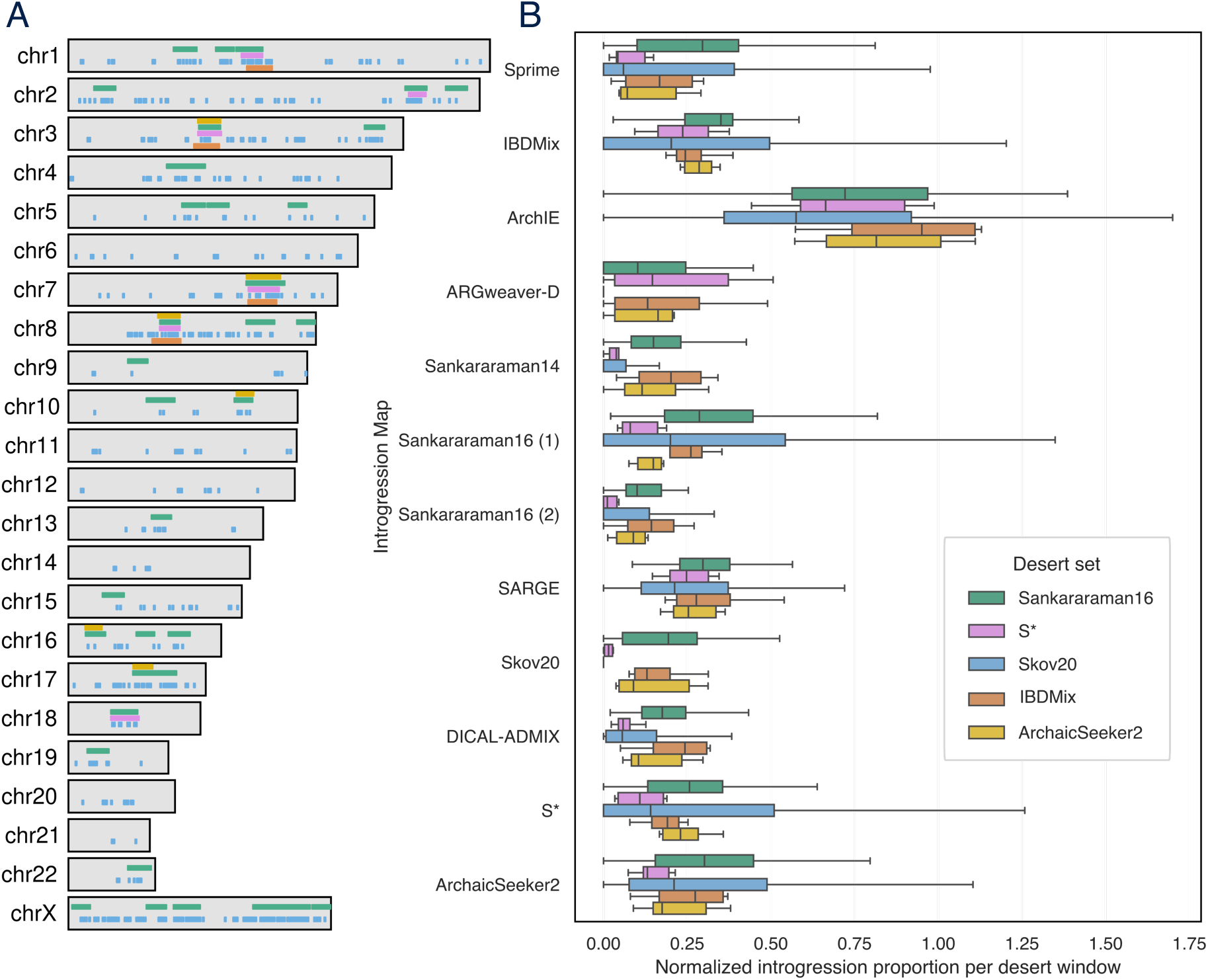
Overlap of introgression deserts across maps. (A) Locations of Neanderthal introgression deserts identified by five previous methods across the genome. (B) Proportion of introgression called in each desert set normalized to each map’s genome-wide introgression proportion. Each box-and-whisker plot represents the distribution of the proportion of Neanderthal introgression across each map in a Neanderthal desert window, divided by that map’s genome-wide introgression proportion.

Given the substantial differences in criteria used to define deserts in previous studies (Supplementary Table 3), we did not expect full agreement between study-specific deserts. Nonetheless, eight distinct genomic regions on chromosomes 3, 7, and 8 harbor deserts identified across all five maps (Figure 8A). Notably, this includes a region on chromosome three that overlaps *ROBO1,* which is involved in neurodevelopment and language abilities^36^, and was previously highlighted in Neanderthal desert analysis^15^. An additional 36 desert regions are supported by at least three maps; these include regions overlapping *FOXP2 and ROBO2*, genes previously proposed to be involved in human-specific traits such as language^8,11,15^. However, genes in these 36 loci are not significantly enriched for any phenotypes using rGREAT.

To evaluate introgression patterns in the annotated deserts across all maps, we compared introgression levels called in each map in desert regions (Figure 8B). For each desert region, we computed the proportion of introgressed bases in the region called by other maps. To account for differences in the overall amount of introgression called by each method, we then normalized each proportion by each method’s genome-wide introgression proportion. Introgression levels were relatively low for many of the deserts, but there were substantial differences between maps. Introgression levels in the Skov20 deserts deviated most from other desert sets, likely due to inferring deserts based on a map of introgression in a large cohort of only modern Finnish individuals. These results suggest several deserts supported by most methods but also highlight substantial variation in what is called an introgression desert dependent on both the method and set of individuals analyzed.

## 3 Discussion

Neanderthal introgression inference methods vary in their use of Neanderthal and outgroup genomes, statistical approaches, and target genomes. Here, we compare a representative set of Neanderthal introgression inference methods and their associated putative introgressed maps in modern humans. We find that support for specific predicted introgressed regions is bimodal, with around half of the regions highly supported by most maps and the remainder specific to one or a few maps. As a result, there is substantial variation in the introgression predictions of different methods, even when applied to the same individual. Focusing on the effects of this variation on downstream analyses, we find that some evolutionary patterns are conserved across all maps, such as evidence for weaker background selection in regions with introgression. However, functional annotations of genes in introgressed regions and the identification of introgression deserts depend on the introgression map used.

We hope that the comparison and synthesis of Neanderthal introgression maps provided here will enable future studies to easily evaluate the robustness of their conclusions to the introgression map used and support the development of more accurate methods for calling introgression. To this end, we suggest that studies focused on a specific introgressed variant or region ensure that it is supported by maps based on different input data, algorithms, and assumptions. If a region is only supported by one method, this does not necessarily mean that it is not introgressed, but it suggests that further locus-specific modeling is required to resolve its history. For analyses that study genome-wide patterns of introgression, we suggest evaluating whether results are qualitatively similar when using multiple different maps and the core set of highly supported regions. At the individual level, we provide introgression maps for 84 1KG CEU individuals that have been used in many previous studies of introgression. Researchers developing new Neanderthal introgression detection algorithms can use these individual-level introgression maps as an additional source of comparison to existing methods. We also provide introgression maps for 1KG and SGDP African genomes when available, and we encourage the further development of algorithms to infer introgression in individuals from Africa.

Our comparison of introgression desert regions called by different introgression maps found that previously highlighted genes, including *ROBO1, ROBO2*, and *FOXP2,* were included in these deserts, with the *ROBO1* desert supported by all five methods. The consistency of the desert regions overlapping these genes supports their potential role in modern human-specific biology, and further investigation into these desert regions may reveal the underpinnings of modern human evolution after Neanderthal introgression. However, we observed that introgression levels predicted in many previously called desert regions vary greatly by method.

Our analyses have several limitations that should be kept in mind when interpreting our conclusions and using the synthesized introgression maps. First, there is no “ground truth” available to evaluate introgression predictions. We do not know the exact history of interactions between Neanderthals and modern humans and how these events influenced our genomes. Moreover, modern human populations have diverse demographic histories that are challenging to infer and integrate into these analyses. Some previous studies, such as ArchIE, have used simulations to create genomes in which the true history of all loci is known, but such approaches are heavily dependent on the assumptions of the simulation. Given the lack of ground truth, we focus on quantifying differences and similarities between introgression maps and are unable to definitively determine whether discordance reflects the inaccuracy of a method.

Second, the set of introgressed regions detected depends on the number and genetic diversity of genomes studied. Given challenges of running some of the methods, we were unable to generate predictions for all methods on the same set of individuals. As a result, we carried out several different analyses, limiting each introgression map to contain the same number of base pairs or subsetting prediction cohorts to the same individuals across methods. In most cases, these analyses revealed similar conclusions to analyzing the entire published maps.

Third, multiple methods and ontologies exist to perform phenotype enrichment on a set of genomic loci, which may yield different enrichment strengths and phenotypes implicated. As an illustrative test case, we quantified enrichment in regions with Neanderthal introgression in modern humans using HPO, which consists of gene annotations inferred primarily from human monogenic disease phenotypes. The enrichments we observe reflect many of the previously reported enrichments for phenotypes relevant to known differences between Neanderthals and humans, such as optical^10^, reproductive^37^, and skeletal^19,24,38^ morphology. While these are present in introgressed regions supported by multiple methods, they are far from consistent across methods. We also note that other annotation sets and enrichment testing methods could identify different phenotype enrichment patterns.

Finally, all our analyses were carried out in the context of the hg19 genome assembly. This was necessary given the focus of previous Neanderthal variant calls and introgression analyses on this assembly. To facilitate future analyses, we provide our integrated prediction sets lifted over to both the hg38 and the CHM13 telomere-to-telomere (T2T) assembly. We anticipate that our findings will hold for these newer assemblies; however, additional introgressed loci would likely be found if the full detection pipelines were applied to these genome assemblies. Recent studies have begun to focus on the complete human reference genome^39^, T2T-CHM13, and we encourage future work on Neanderthal introgression to focus on this assembly.

Given our limited understanding of the legacy of Neanderthals on humans today and the continued discovery of archaic genomes, we anticipate further development of algorithms to identify introgressed DNA and new applications for these methods. We hope that our results will guide these studies and ensure robust conclusions. Additionally, introgressed DNA identified between methods from other ancient hominin species, such as Denisovans, has yet to be formally evaluated. By quantifying differences in introgression inference methods from archaic species, we will gain a greater understanding of how humans evolved and how our ancestors shaped our history.

## 4 Methods

### 4.1 Defining and harmonizing introgression maps

For each introgression inference method, we obtained the predicted introgressed map from the original study. Each introgression map was then processed in a study-specific manner to generate standardized human genome build hg19-referenced bed files consisting of introgressed regions and an associated confidence score. For each method, introgressed fragments were merged across population groupings using BEDtools^40^ v2.31.1 merge. The highest scores across merged regions were retained. To generate method-specific introgression maps, all detected introgressed loci across populations were merged. All introgression maps were built in hg19 due to its overwhelming use in ancient DNA studies.

**Chen 2020 (IBDMix):** Introgressed segments were downloaded from https://drive.google.com/drive/folders/1mDQaDFS-j22Eim5_y7LAsTTNt5GWsoow, as described in the original IBDMix paper^15^. IBDMix uses LOD scores, reflecting the likelihood that a region is shared with the reference Neanderthal through identity by descent. Here, all introgressed loci have LOD scores greater than 3, as provided by the original authors. Ancestral groupings are based on 1KG, as well as SGDP for genomes from Papua New Guinea.

**Sankararaman 2014:** Neanderthal introgressed regions and SNPs were downloaded from https://drive.google.com/drive/folders/1jRymYI3GyNrBXKtd6pT9pqoikZ6I8P3Z. Each entry is associated with scores reflecting the probability of introgression from the CRF. All haplotypes have a probability greater than 0.9, as provided in the downloaded data. Ancestral groupings are based on 1KG categorizations.

**Sankararaman 2016:** Introgression maps were downloaded from https://drive.google.com/drive/folders/1DyhMw0E1mXQUDNeGQbQrcQbHe0uTyK2h. Introgressed regions were processed separately for two directories available: The first directory only uses African individuals as a non-Neanderthal introgressed outgroup, whereas the second directory uses both African and the Denisovan genome as a non-Neanderthal introgressed outgroup, which is the focus of the 2016 paper. We refer to these two methods, respectively, as Sankararaman16 (1) and Sankararaman16 (2). For each directory, the average predicted probability for each haplotype was obtained to generate introgression maps. All CRF haplotypes have introgression probability greater than 0.9, and variants have Neanderthal ancestry probability in derived alleles greater than 0.5 individuals. Additionally, variant information was generated by separating the SNP ID into chromosome and position. Finally, the “start” column was added by subtracting one from the variant position column.

**Skov 2020:** Supplementary dataset 1 from the original publication^13^ was downloaded, containing all introgressed fragments with an introgression probability greater than 0.9. Using LiftOver^41^, we converted the genomic coordinates to hg19 from hg38 to allow for direct comparison with other methods. To focus on Neanderthal introgressed regions, fragments with an archaic label of Altai or Vindija were subsetted.

**SARGE (Schaefer 2021):** Data were provided by Dr. Nathan Schaefer, by request. Data provided were already in the hs37d5 genome build. Scores reflect similarity scores between haplotypes, depending on how many ancestral recombination events are shared, and all data provided are thresholded according to the original implementation^17^.

**Browning 2018 (Sprime):** Sprime output for 1KG (non-African) and SGDP Papuan individuals were downloaded from https://doi.org/10.17632/y7hyt83vxr.1, consisting of variant-level and fragment-level files. For variant files, a “start” column was added by subtracting one from the variant position column. Fragment files were generated by taking the first and last variant positions for each Segment_ID. To subset for Neanderthal-specific introgressed data, we subsetted Sprime output where NMATCH contained “match.” Sprime scores reflect the confidence of introgression associated with each loci, and all Sprime loci met a minimum threshold value of 150,000.

**Vernot 2016 (S*):** S* Neanderthal haplotypes and variants were downloaded from https://drive.google.com/drive/folders/0B9Pc7_zItMCVWUp6bWtXc2xJVkk?resourceke <u>y=0-Cj8G4QYndXQLVIGPoWKUjQ</u>. Neanderthal-specific population and chromosome-specific merged files in the introgressed_haplotypes folder were used to generate Neanderthal introgression maps. Scores reflect the posterior probability Neanderthal introgression for each haplotype, labelled post.p.alt1 in the provided data. Variants were taken from introgressed_tag_snp_frequencies.

**Hubisz 2020 (ARGweaver-D):** Introgressed haplotypes from Argweaver-D were obtained from http://compgen.cshl.edu/ARGweaver/introgressionHub/. Introgressed regions from the ooaM1A directory, reflecting a model with migration bands from Neanderthals and Denisovans into 4 modern humans from SGDP: 2 Africans (Mandenka and Khomani San), one Papuan, and one Basque individual. Files used to generate a combined file were ooaM1A/Papuan_1F.bed, ooaM1A/Basque_2F.bed, ooaM1A/Mandenka_2F.bed, and ooaM1A/Khomani_San_1F.bed. All loci labelled as neaTo(Human) were included, and scores represent heterozygous (500) vs homozygous (1000) introgressed regions.

**ArchIE (Durvasula 2019):** Neanderthal introgressed haplotypes in CEU 1KG individuals were provided by Dr. Arun Durvasula, by request. Haplotypes with probability of Neanderthal introgression greater than 0.9984 were subsetted, reflecting ∼2% Neanderthal introgression per individual. To avoid overlapping 1 base pair windows, we subtracted 1 from each end coordinate.

**ArchaicSeeker2 (Yuan 2021):** Introgressed fragments for 1KG and SGDP Papuan individuals were obtained from .seg files in the IntrogressedSeg directory at https://github.com/Shuhua-Group/ArchaicSeeker2.0. Segments with “Neanderthal” for the BestMatchedPop columns were retained to generate introgression maps.

**DICAL-ADMIX (Steinrücken 2018):** DICAL-ADMIX output for CEU, CHB, and CHS 1KG individuals were downloaded from https://dical-admix.sourceforge.net. All fragments provided have posterior probabilities of Neanderthal introgression greater than 0.42 and were used to generate introgression maps.

### 4.2 Jaccard similarity and hierarchical clustering of introgression maps

Jaccard statistics were computed by dividing the number of shared base pairs over the union of base pairs between pairs of methods !, # with NumPy v1.26.3. Normalized Jaccard similarity was computed by using the smaller set of base pairs as the denominator, rather than the union between two sets.

Hierarchical clustering was performed using the linkage function in the SciPy v1.14.1 on the raw jaccard distance matrix, computed by subtracting each Jaccard similarity value from 1. Clustering was performed using the Farthest Point Algorithm, where $(!, #) = max ($,-.(![,], #[1]).

To obtain the top 793.9 Mb of introgressed loci for each method, introgressed regions for each introgression map were sorted by highest associated scores and subsetted.

Results were visualized using upsetplot v0.9.0, seaborn v0.13.2, and matplotlib v3.9.2 in Python v3.11.5.

### 4.3 B-statistic scores in introgressed versus non-introgressed regions

B statistic scores from McVicker et al^34^ were obtained from Telis et al^27^, containing B statistic values ranging from 1-1,000. B statistic values were quantized into multiples of 50, and lifted to hg19 by Telis et al^27^. To compute B statistic values across introgression maps, we assigned each 500 bp window across the hg19 genome as either introgressed or non-introgressed. A window was introgressed if it contained at least one introgressed region, as we hoped to observe whether windows with tolerance for some introgression had different B statistic scores compared to non-introgressed windows.

Next, we performed bootstrapping with 1,000 iterations to obtain a bootstrapped mean, along with confidence intervals. We visualized results using matplotlib v3.9.2, and computed differences between B statistic values between introgressed and non-introgressed windows using a Mann-Whitney U test.

### 4.4 Phenotype annotation enrichment in introgression maps

Gene set enrichment analysis for each introgression map was performed using rGREAT v2.6.0^42^ with the default hg19 genome (gaps removed) as the enrichment background and gene sets in the Human Phenotype Ontology, which we selected to associate loci with interpretable human features. Each gene in the ontology was extended to capture short and long-range transcription start site (TSS) associations using default parameters (basal domain and extension around the transcription start site, 5 kb upstream and 1 kb downstream, then extension of the basal domain in both directions to 1 Mb, or until the neighbor gene’s basal domain is reached to capture long-range associations). Next, rGREAT computes the fraction of genome overlap between the set of input regions and gene-associated extended regions for each phenotype’s gene set, then computes an enrichment p-value under a binomial model.

Additionally, we used EnrichR v3.4^43^ as a complimentary gene set enrichment method for further comparison. To obtain an input gene set consisting of genes that overlapped with exonic regions for a set of input loci, we used Annotatr v.1.30.0^44^. We annotated our input bed file with hg19 genes where an input region falls in an exon (hg19_genes_exons), and annotations were built by Annotatr’s *build_annotations* function using data from the *TxDb.Hsapiens.UCSC.hg19* database. The exon-specific gene list from annotatR was used as an input for EnrichR, and gene set libraries were limited to GWAS Catalog 2023 and HPO.

Principal components analysis (PCA) was performed in R v4.4.3 on adjusted p-values from EnrichR and rGREAT with the prcomp function from stats v4.4.3. Adjusted p-values were standardized, and phenotypes with constant or NA values were excluded, resulting in 386 phenotypes included in the PCA.

Visualizations were generated using ggplot2 v3.5.1 and UpSetR v1.4.0.

### 4.5 Comparing Neanderthal introgression deserts

Neanderthal introgression deserts were obtained for each respective study. IBDMix and S* deserts were manually entered into bed format from Table S8^15^. ArchaicSeeker2 deserts were obtained from Supplementary Table 6^25^, and Skov20 deserts were downloaded from Supplementary Dataset 4^13^ and lifted to hg19 from hg38. Sankararaman16 deserts were provided by Dr. Sriram Sankararaman. No additional modifications to these data were performed.

To observe the amount of introgression for each desert window, we used BEDtools coverage to compute the fraction of introgressed loci for each method’s introgression map, previously described, overlapping each desert window.

Desert regions across the genome for each method were plotted using karyoploteR^45^.

## 4.6 Data availability

The publicly available data used for analysis are available in the following repositories: https://github.com/yaenchen/NeanderthalIntrogressionMaps.

## 4.7 Code availability

The publicly available code for analysis is available in the following repositories: https://github.com/yaenchen/NeanderthalIntrogressionMaps.

## Acknowledgements

We thank members of the Capra Lab for their insight, guidance, and support on this work. This work was supported by the National Institutes of Health (NIH) General Medical Sciences award R35GM127087 to JAC. KAV was supported by NIH award T32CA108462. This work was performed on the Wynton high-performance compute cluster, which is supported by UCSF research faculty and UCSF institutional funds. The authors wish to thank the UCSF Wynton team for their ongoing technical support of the Wynton environment.

## Author Contributions

Yaen Chen: Conceptualization, Methodology, Software, Formal analysis, Investigation, Data curation, Writing – Original Draft, Writing – Review & Editing, Visualization. Keila Velazquez-Arcelay: Data Curation, Writing – Review & Editing. John A. Capra: Conceptualization, methodology, investigation, Writing – Original Draft, Writing – Review & Editing, Supervision, Funding Acquisition.

## References

1. Higham, T. et al. The timing and spatiotemporal patterning of Neanderthal disappearance. Nature 512, 306–309 (2014).

2. Mafessoni, F. et al. A high-coverage Neandertal genome from Chagyrskaya Cave. Proceedings of the National Academy of Sciences 117, 15132–15136 (2020).

3. Prüfer, K. et al. A high-coverage Neandertal genome from Vindija Cave in Croatia. Science 358, 655–658 (2017).

4. Prüfer, K. et al. The complete genome sequence of a Neanderthal from the Altai Mountains. Nature 505, 43–49 (2014).

5. Ahlquist, K. D. et al. Our Tangled Family Tree: New Genomic Methods Offer Insight into the Legacy of Archaic Admixture. Genome Biology and Evolution 13, evab115 (2021).

6. Green, R. E. et al. A Draft Sequence of the Neandertal Genome. Science 328, 710–722 (2010).

7. Iasi, L. N. M. et al. Neandertal ancestry through time: Insights from genomes of ancient and present-day humans. bioRxiv 2024.05.13.593955 (2024) doi:10.1101/2024.05.13.593955.

8. Vernot, B. et al. Excavating Neandertal and Denisovan DNA from the genomes of Melanesian individuals. Science 352, 235–239 (2016).

9. Steinrücken, M., Spence, J. P., Kamm, J. A., Wieczorek, E. & Song, Y. S. Model-based detection and analysis of introgressed Neanderthal ancestry in modern humans. Molecular Ecology 27, 3873–3888 (2018).

10. Sankararaman, S. et al. The genomic landscape of Neanderthal ancestry in present-day humans. Nature 507, 354–357 (2014).

11. Sankararaman, S., Mallick, S., Patterson, N. & Reich, D. The Combined Landscape of Denisovan and Neanderthal Ancestry in Present-Day Humans. Current Biology 26, 1241–1247 (2016).

12. Skov, L. et al. Detecting archaic introgression using an unadmixed outgroup. PLOS Genetics 14, e1007641 (2018).

13. Skov, L. et al. The nature of Neanderthal introgression revealed by 27,566 Icelandic genomes. Nature 582, 78–83 (2020).

14. Browning, S. R., Browning, B. L., Zhou, Y., Tucci, S. & Akey, J. M. Analysis of Human Sequence Data Reveals Two Pulses of Archaic Denisovan Admixture. Cell 173, 53–61.e9 (2018).

15. Chen, L., Wolf, A. B., Fu, W., Li, L. & Akey, J. M. Identifying and Interpreting Apparent Neanderthal Ancestry in African Individuals. Cell 180, 677–687.e16 (2020).

16. Hubisz, M. J., Williams, A. L. & Siepel, A. Mapping gene flow between ancient hominins through demography-aware inference of the ancestral recombination graph. PLOS Genetics 16, e1008895 (2020).

17. Schaefer, N. K., Shapiro, B. & Green, R. E. An ancestral recombination graph of human, Neanderthal, and Denisovan genomes. Science Advances 7, eabc0776.

18. Durvasula, A. & Sankararaman, S. A statistical model for reference-free inference of archaic local ancestry. PLOS Genetics 15, e1008175 (2019).

19. McArthur, E., Rinker, D. C. & Capra, J. A. Quantifying the contribution of Neanderthal introgression to the heritability of complex traits. Nature Communications 12, 4481 (2021).

20. Velazquez-Arcelay, K. et al. Archaic Introgression Shaped Human Circadian Traits. Genome Biology and Evolution 15, evad203 (2023).

21. Zeberg, H. & Pääbo, S. The major genetic risk factor for severe COVID-19 is inherited from Neanderthals. Nature 587, 610–612 (2020).

22. Koller, D. et al. Denisovan and Neanderthal archaic introgression differentially impacted the genetics of complex traits in modern populations. BMC Biology 20, 249 (2022).

23. Dannemann, M. et al. Neandertal introgression partitions the genetic landscape of neuropsychiatric disorders and associated behavioral phenotypes. Translational Psychiatry 12, 433 (2022).

24. Wei, X. et al. The lingering effects of Neanderthal introgression on human complex traits. eLife 12, e80757 (2023).

25. Yuan, K. et al. Refining models of archaic admixture in Eurasia with ArchaicSeeker 2.0. Nature Communications 12, 6232 (2021).

26. McCoy, R. C., Wakefield, J. & Akey, J. M. Impacts of Neanderthal-Introgressed Sequences on the Landscape of Human Gene Expression. Cell 168, 916–927.e12 (2017).

27. Telis, N., Aguilar, R. & Harris, K. Selection against archaic hominin genetic variation in regulatory regions. Nature Ecology & Evolution 4, 1558–1566 (2020).

28. Petr, M., Pääbo, S., Kelso, J. & Vernot, B. Limits of long-term selection against Neandertal introgression. Proceedings of the National Academy of Sciences 116, 1639–1644 (2019).

29. Zhang, X. et al. Neanderthal introgressed ancestry reveals human genomic regions enriched with recessive deleterious mutations. bioRxiv 2025.05.07.652751 (2025) doi:10.1101/2025.05.07.652751.

30. Harris, K. & Nielsen, R. The Genetic Cost of Neanderthal Introgression. Genetics 203, 881–891 (2016).

31. Reilly, P. F., Tjahjadi, A., Miller, S. L., Akey, J. M. & Tucci, S. The contribution of Neanderthal introgression to modern human traits. Current Biology 32, R970–R983 (2022).

32. Auton, A. et al. A global reference for human genetic variation. Nature 526, 68–74 (2015).

33. Kerdoncuff, E. et al. 50,000 years of Evolutionary History of India: Insights from ∼2,700 Whole Genome Sequences. bioRxiv 2024.02.15.580575 (2024) doi:10.1101/2024.02.15.580575.

34. McVicker, G., Gordon, D., Davis, C. & Green, P. Widespread Genomic Signatures of Natural Selection in Hominid Evolution. PLOS Genetics 5, e1000471 (2009).

35. Racimo, F., Sankararaman, S., Nielsen, R. & Huerta-Sánchez, E. Evidence for archaic adaptive introgression in humans. Nature Reviews Genetics 16, 359–371 (2015).

36. Hannula-Jouppi, K. et al. The Axon Guidance Receptor Gene ROBO1 Is a Candidate Gene for Developmental Dyslexia. PLOS Genetics 1, e50 (2005).

37. Weaver, T. D. & Hublin, J.-J. Neandertal birth canal shape and the evolution of human childbirth. Proceedings of the National Academy of Sciences 106, 8151–8156 (2009).

38. Colbran, L. L. et al. Inferred divergent gene regulation in archaic hominins reveals potential phenotypic differences. Nature Ecology & Evolution 3, 1598–1606 (2019).

39. Liang, S.-A. et al. A refined analysis of Neanderthal-introgressed sequences in modern humans with a complete reference genome. Genome Biology 26, 32 (2025).

40. Quinlan, A. R. & Hall, I. M. BEDTools: a flexible suite of utilities for comparing genomic features. Bioinformatics 26, 841–842 (2010).

41. Perez, G. et al. The UCSC Genome Browser database: 2025 update. Nucleic Acids Research 53, D1243–D1249 (2025).

42. Gu, Z. & Hübschmann, D. rGREAT: an R/bioconductor package for functional enrichment on genomic regions. Bioinformatics 39, btac745 (2023).

43. Chen, E. Y. et al. Enrichr: interactive and collaborative HTML5 gene list enrichment analysis tool. BMC Bioinformatics 14, 128 (2013).

44. Cavalcante, R. G. & Sartor, M. A. annotatr: genomic regions in context. Bioinformatics 33, 2381–2383 (2017).

45. Gel, B. & Serra, E. karyoploteR: an R/Bioconductor package to plot customizable genomes displaying arbitrary data. Bioinformatics 33, 3088–3090 (2017).

