## Supplementary Figures and Tables for "Comparing Neanderthal introgression maps reveals core agreement but substantial heterogeneity"

### Supplementary Information

#### Supplemental Figures and Tables

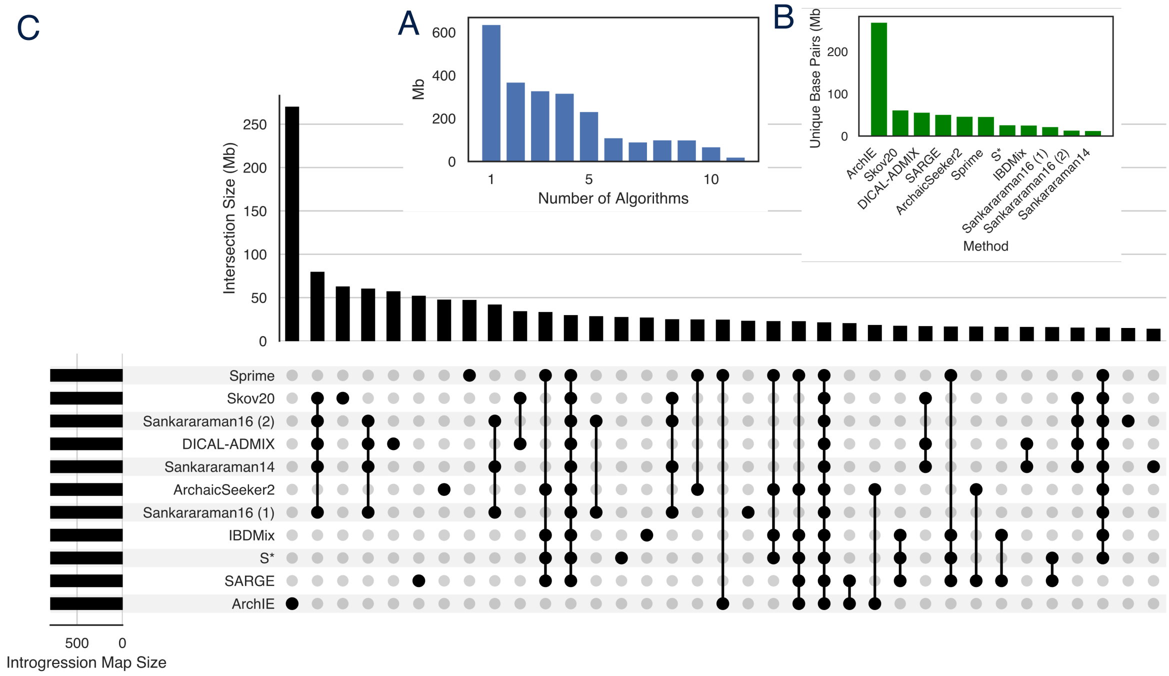


**Supplementary Figure 1**: To account for differences in the total amount of introgression predicted, this comparison is based on the top 793.9 Mb from each map. (A) Histogram of the number of introgression maps supporting each bp predicted as introgressed in at least one map, for size-matched maps. (B) The number of unique base pairs predicted as introgressed by each size-matched map. (C) Upset plot of the patterns of overlap among regions across size-matched introgression maps.

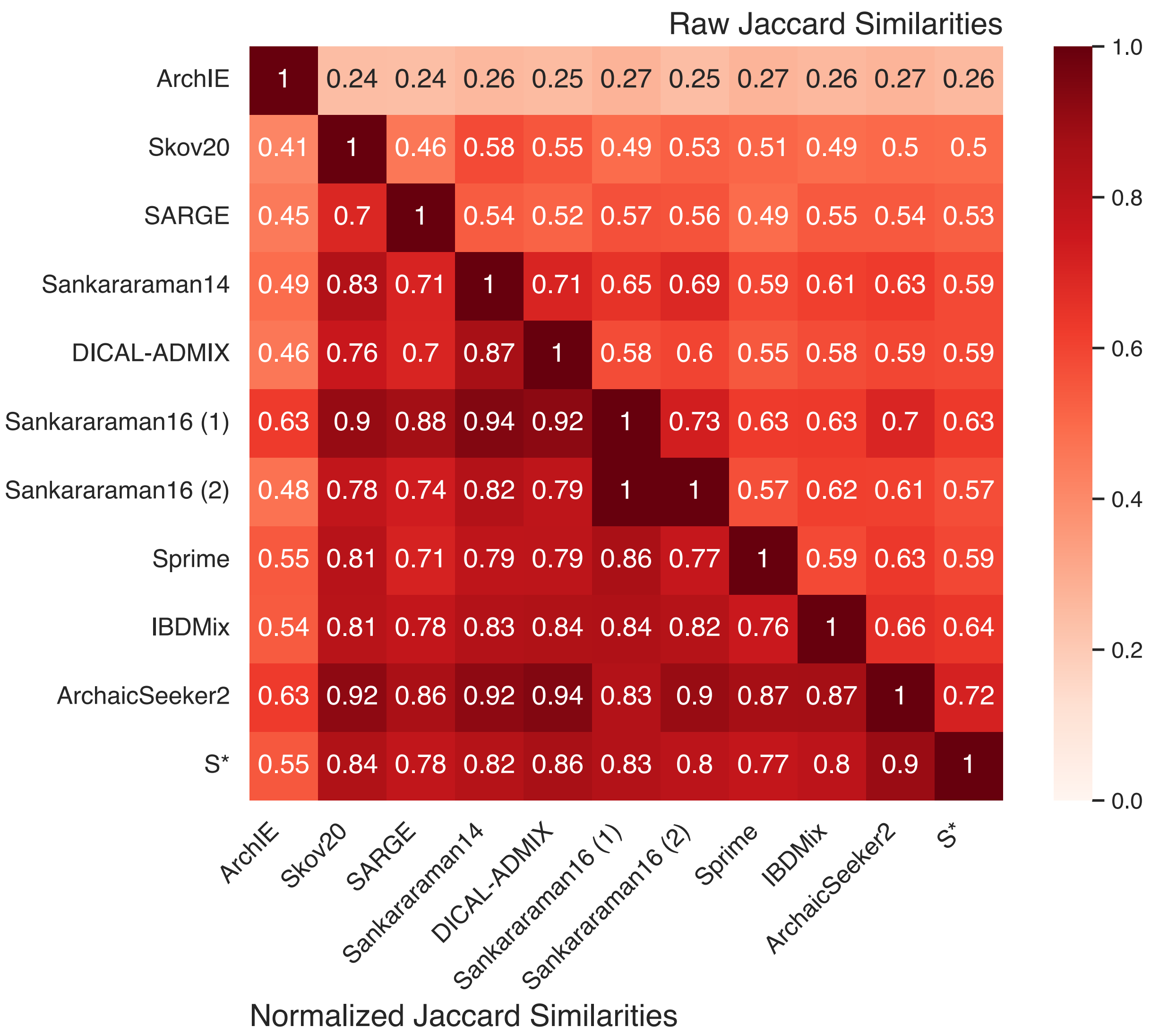


**Supplementary Figure 2:** Raw and normalized Jaccard across complete introgression maps, autosomes only.

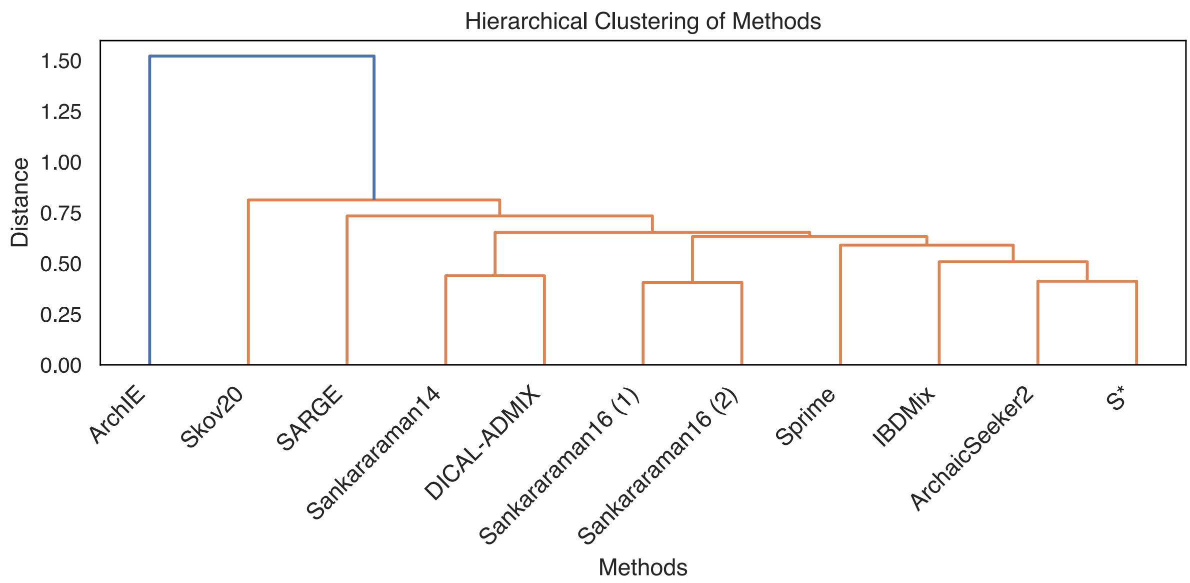


**Supplementary Figure 3: Hierarchical clustering between methods based on Jaccard distances from Supplementary Figure 1.**

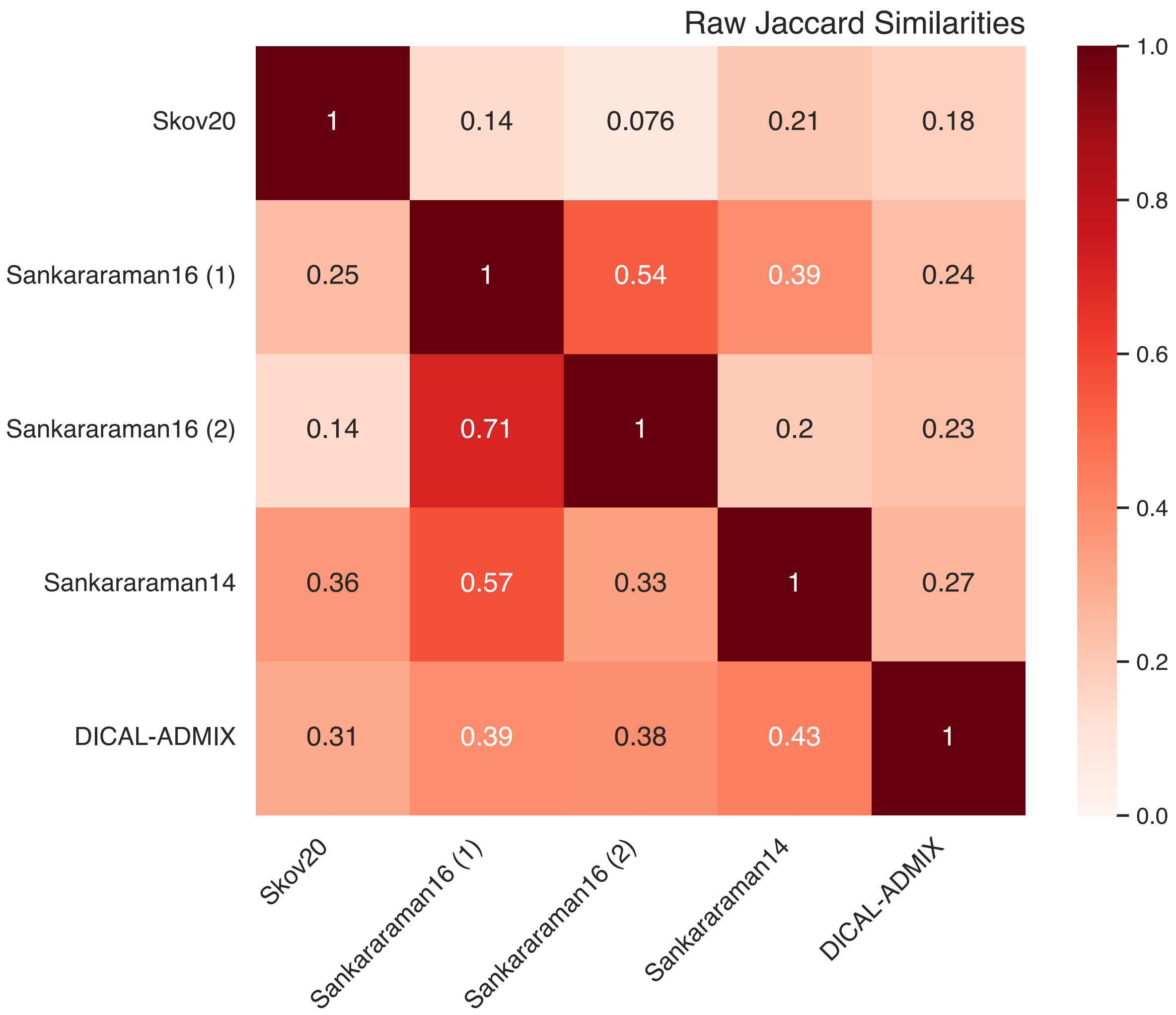


Normalized Jaccard Similarities

**Supplementary Figure 4: Raw and normalized Jaccard across the X chromosome, subsetted to top 7,814,066 bases to match Skov20.**

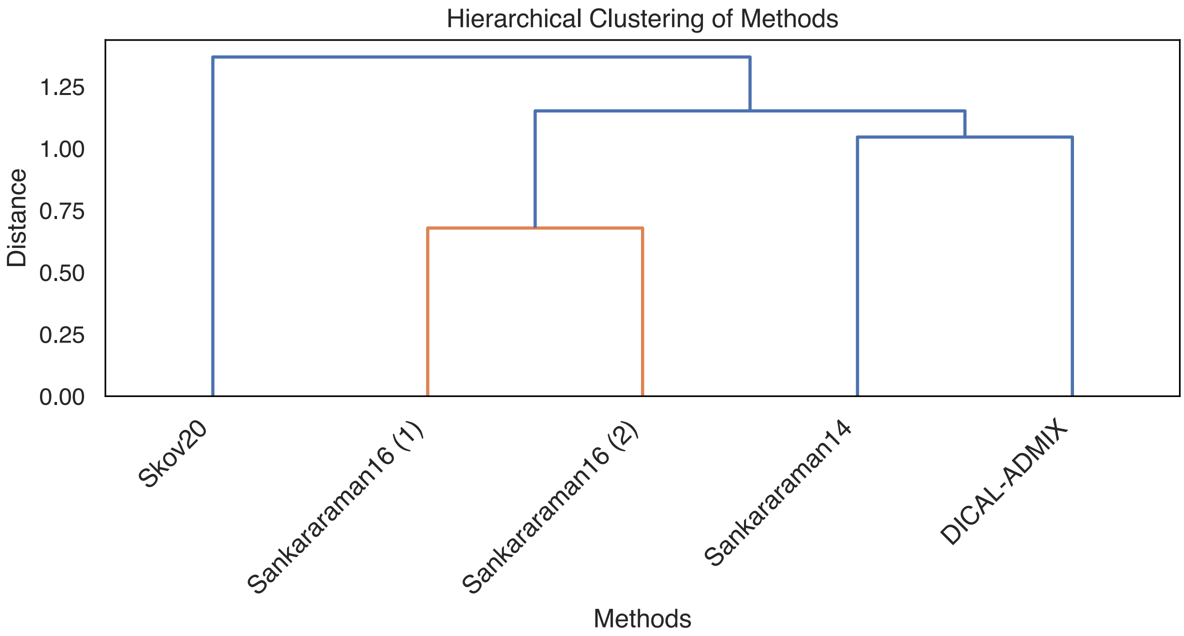


**Supplementary Figure 5: Hierarchical clustering between methods based on Jaccard distances from Supplementary Figure 3.**

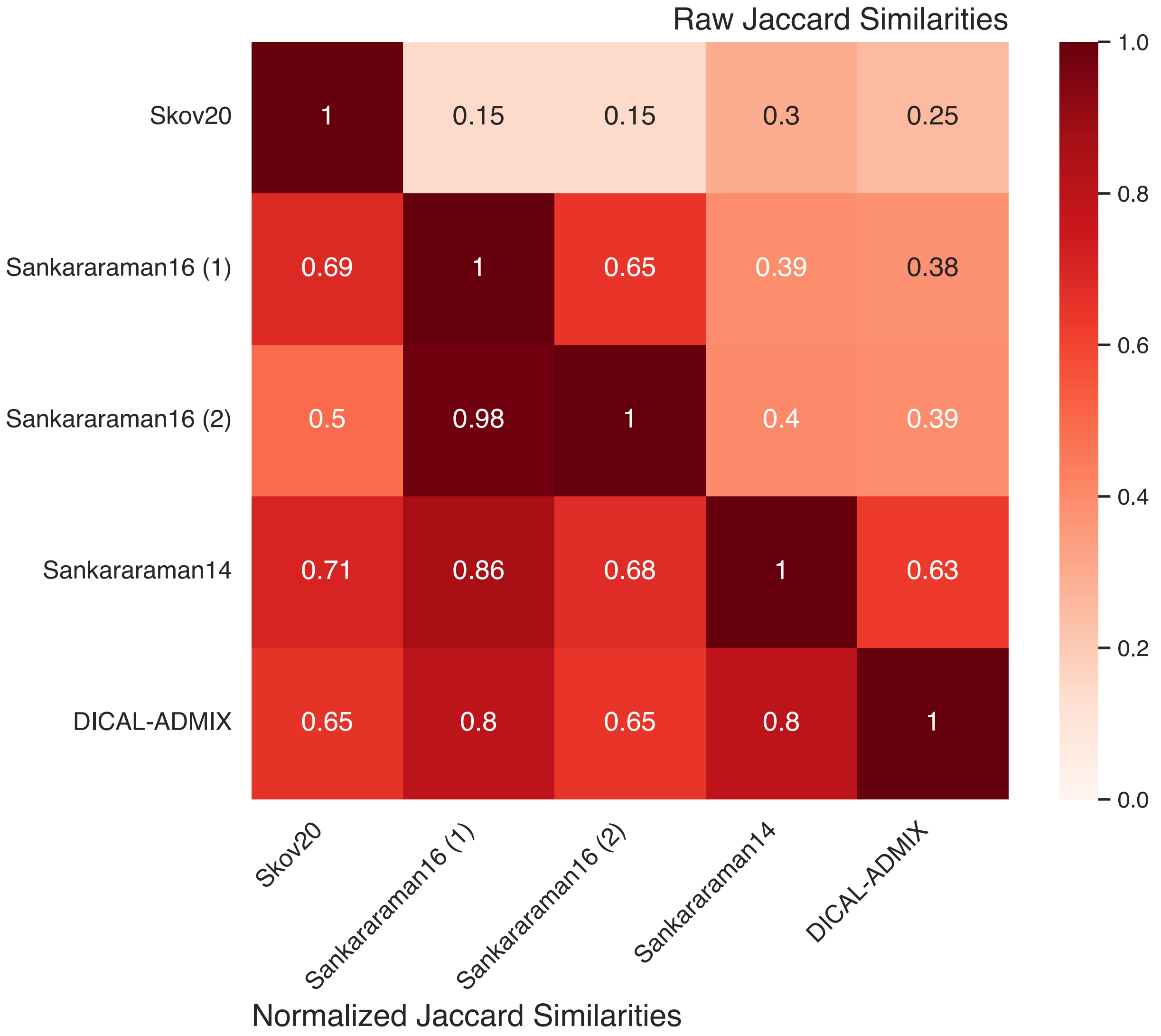


**Supplementary Figure 6: Raw and normalized Jaccard across complete introgression maps, X chromosome only.**

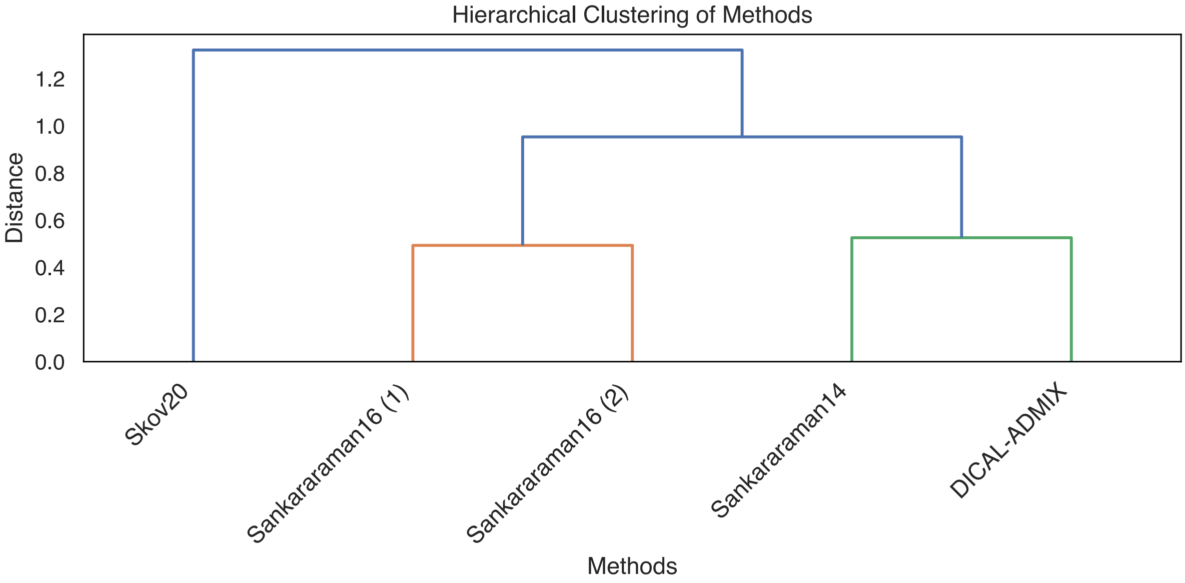

**Supplementary Figure 7: Hierarchical clustering between methods based on Jaccard distances from Supplementary Figure 5.**

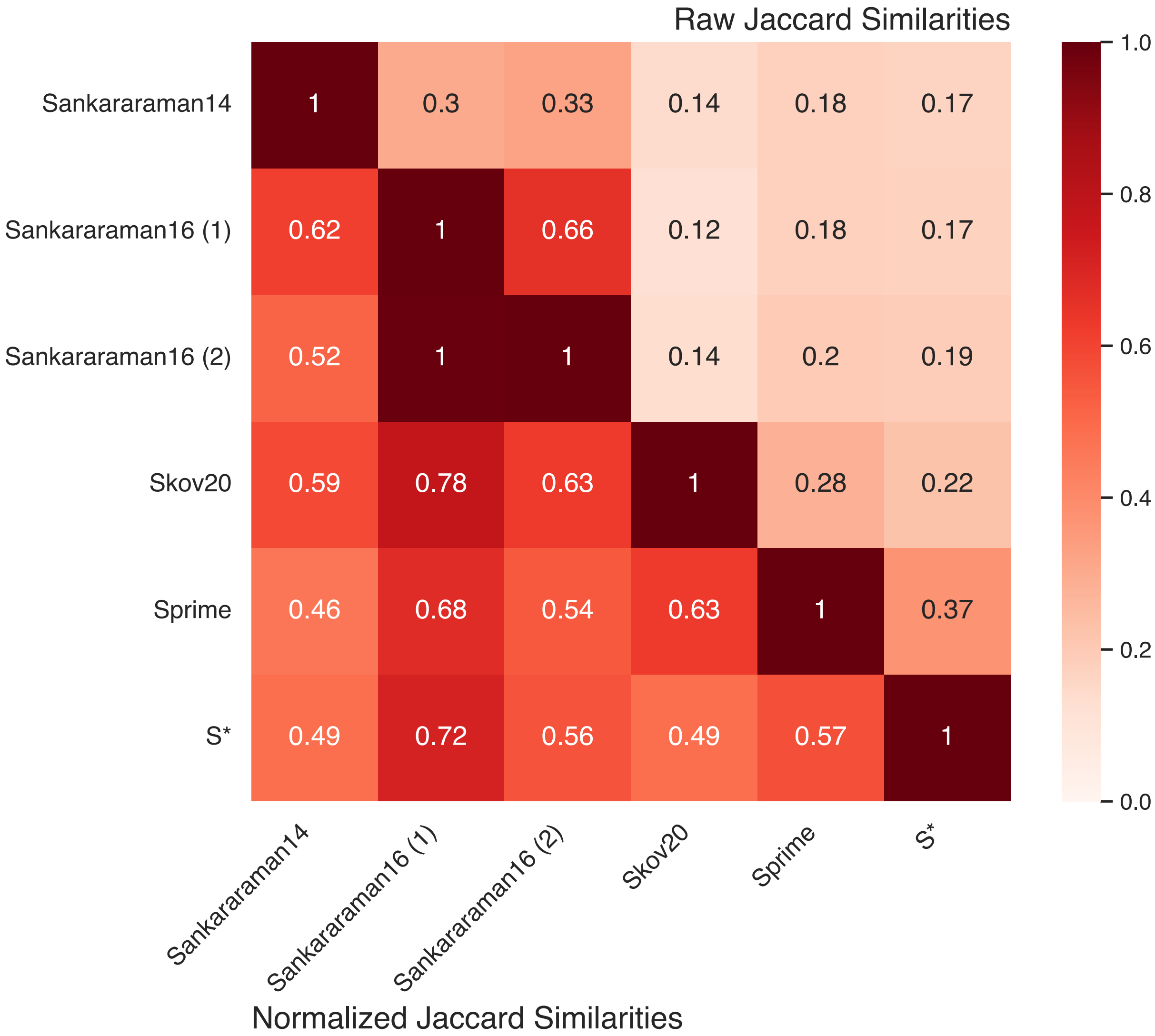


**Supplementary Figure 8: Raw and normalized Jaccard across variants, autosomes only.**

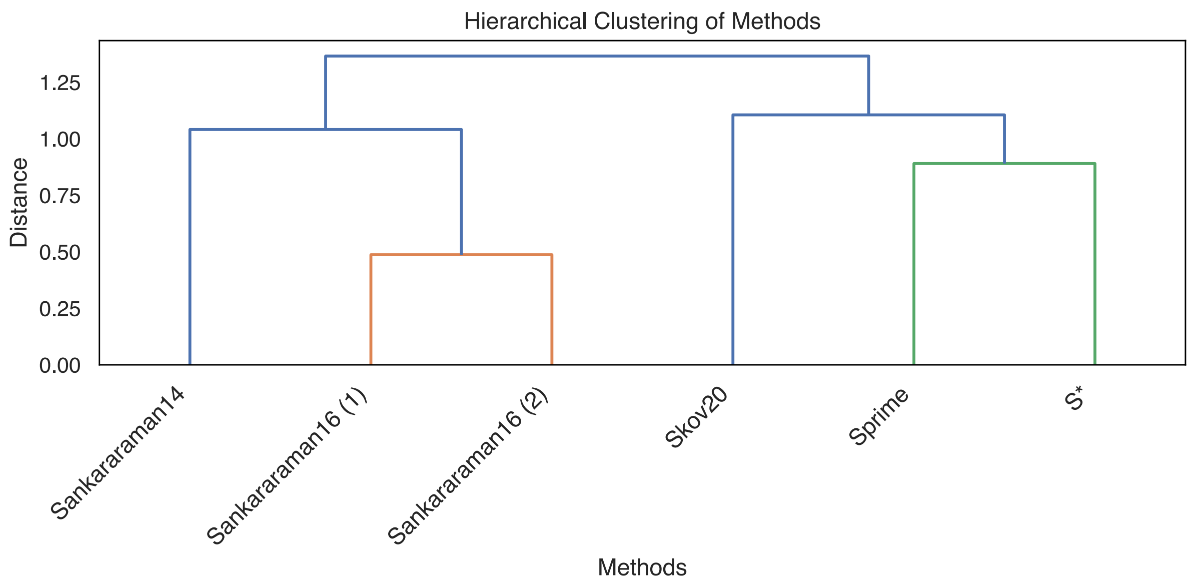


**Supplementary Figure 9: Hierarchical clustering between methods based on Jaccard distances from Supplementary Figure 7.**

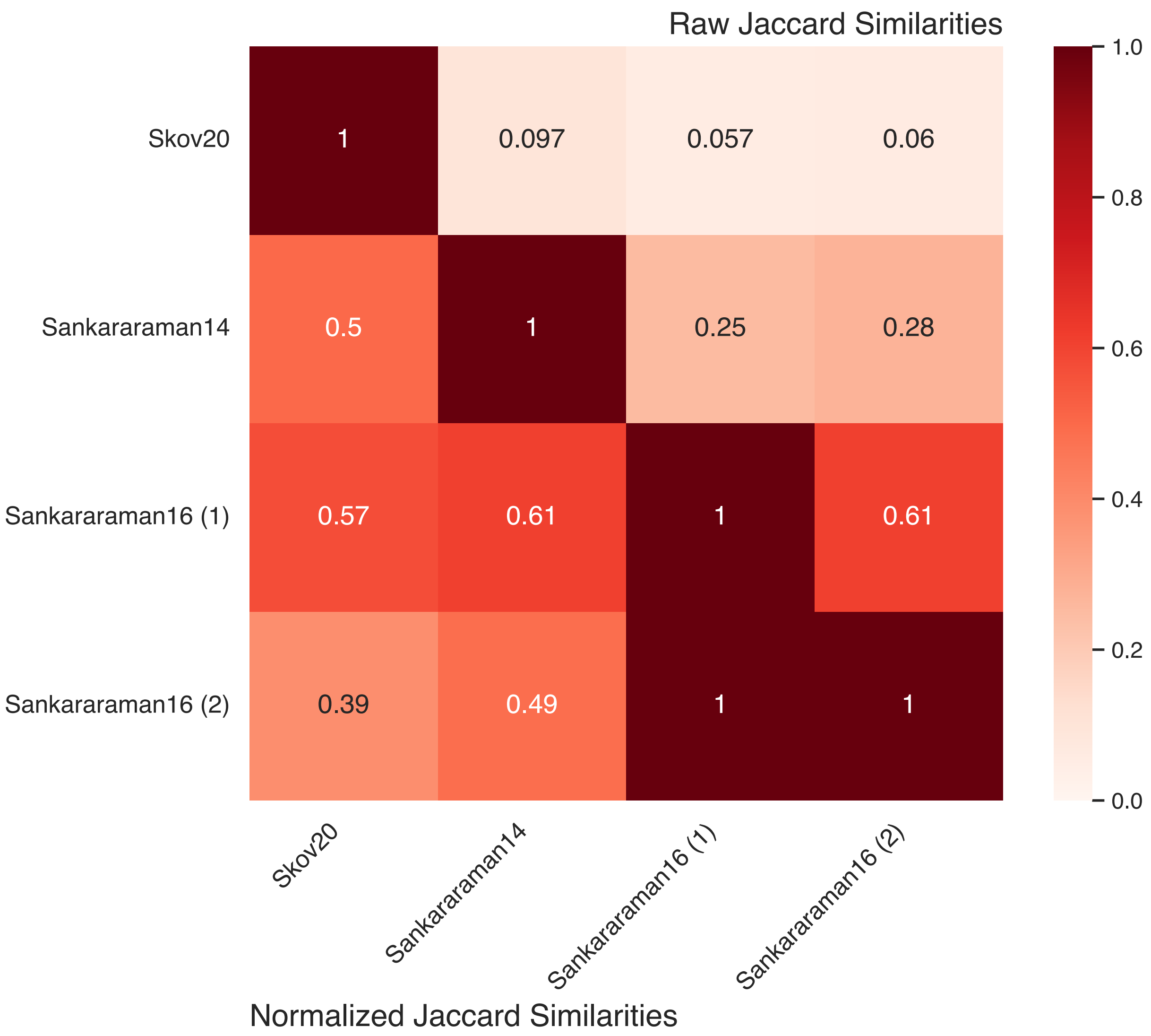


**Supplementary Figure 10: Raw and normalized Jaccard across variants, X chromosome only.**

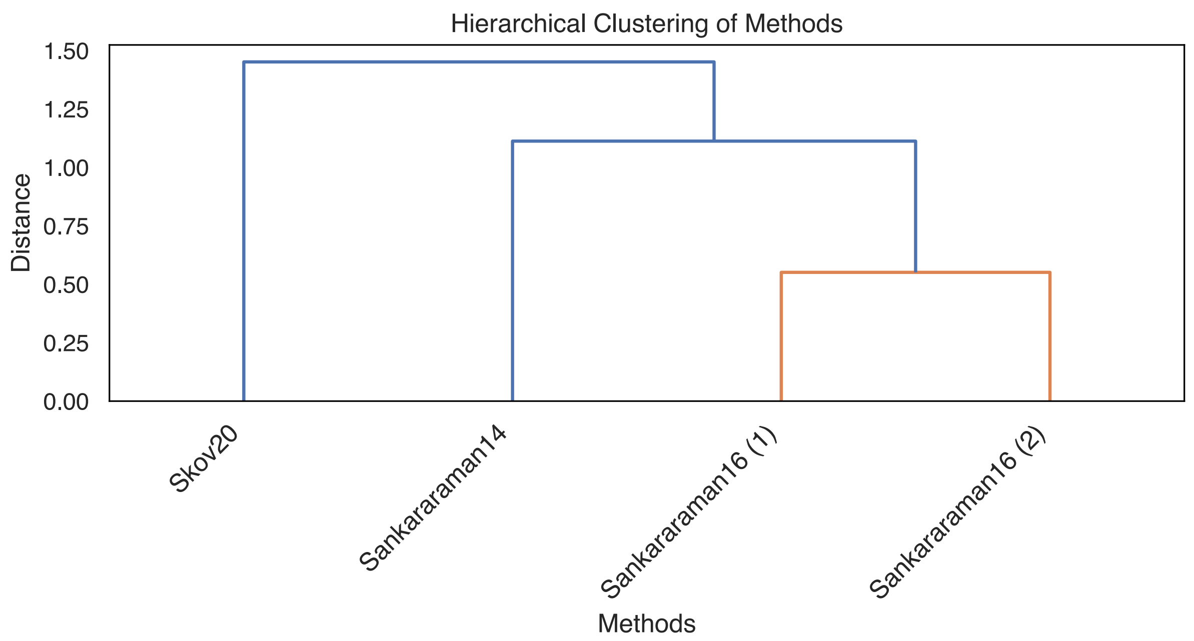


**Supplementary Figure 11: Hierarchical clustering between methods based on Jaccard distances from Supplementary Figure 9.**

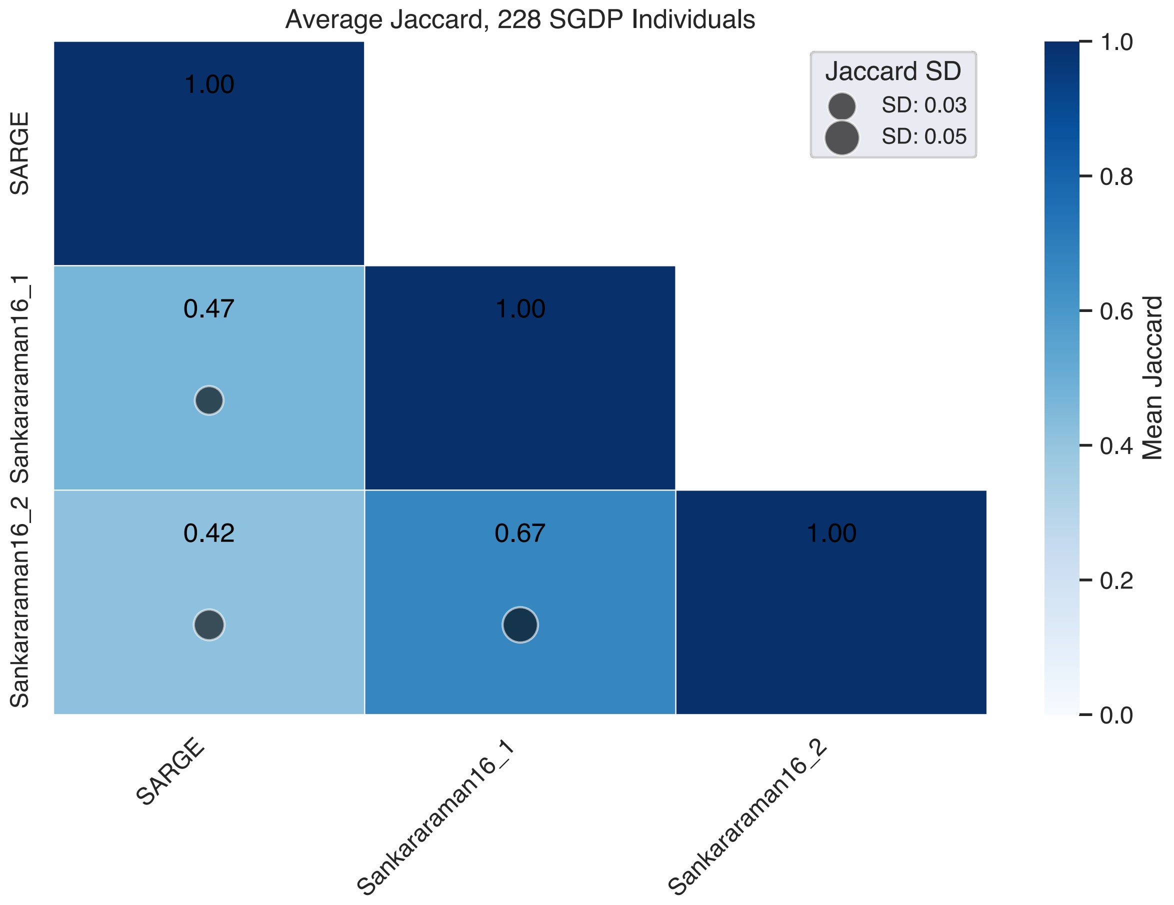

**Supplementary Figure 12:** Raw and normalized Jaccard across introgressed regions in 228 shared SGDP individuals. For this comparison, introgression maps included SARGE, Sankararaman 2016 (1), and Sankararaman 2016 (2). Individuals represented include: Abkhasian-1, Abkhasian-2, Adygei-1, Adygei-2, Albanian-1, Aleut-1, Aleut-2, Altaian-1, Ami-1, Ami-2, Armenian-1, Armenian-2, Atayal-1, Australian-3, Australian-4, Balochi-1, Basque-1, Basque-2, BedouinB-1, BedouinB-2, Bengali-1, Bengali-2, Bergamo-1, Bergamo-2, Bougainville-1, Bougainville-2, Brahmin-1, Brahmin-2, Brahui-1, Brahui-2, Bulgarian-1, Bulgarian-2, Burmese-1, Burmese-2, Burusho-1, Burusho-2, Cambodian-1, Cambodian-2, Chane-1, Chechen-1, Chukchi-1, Crete-1, Crete-2, Czech-2, Dai-1, Dai-2, Dai-3, Dai-4, Daur-1, Daur-2, Druze-1, Druze-2, Dusun-1, Dusun-2, English-1, English-2, Eskimo, Estonian-1, Estonian-2, Even-1, Even-2, Even-3, Finnish-1, Finnish-2, Finnish-3, French-1, French-2, French-3, Georgian-1, Georgian-2, Greek-1, Greek-2, Han-1, Han-2, Han-3, Hawaiian-1, Hazara-1, Hazara-2, Hezhen-1, Hezhen-2, Hungarian-1, Hungarian-2, Icelandic-1, Icelandic-2, Igorot-1, Igorot-2, Iranian-1, Iranian-2, Iraqi, Irula-1, Irula-2, Itelman-1, Japanese-1, Japanese-2, Japanese-3, Jordanian-1, Jordanian-2, Jordanian-3, Kalash-1, Kalash-2, Kapu-1, Kapu-2, Karitiana-1, Karitiana-2, Karitiana-3, Khonda, Kinh-1, Kinh-2, Korean-1, Korean-2, Kusunda-1, Kusunda-2, Kyrgyz-1, Kyrgyz-2, Lahu-1, Lahu-2, Lezgin-1, Lezgin-2, Madiga-1, Madiga-2, Makrani-1, Makrani-2, Mala-2, Mala-3, Mansi-1, Mansi-2, Maori-1, Mayan-1, Mayan-2, Miao-1, Miao-2, Mixe-1, Mixe-2, Mixe-3, Mixtec-1, Mixtec-2, Mongola-1, Mongola-2, Naxi-1, Naxi-2, Naxi-3, North, Norwegian-1, Orcadian-1, Orcadian-2, Oroqen-1, Oroqen-2, Palestinian-1, Palestinian-2, Palestinian-3, Papuan-1, Papuan-10, Papuan-11, Papuan-12, Papuan-13, Papuan-14, Papuan-15, Papuan-2, Papuan-3, Papuan-4, Papuan-5, Papuan-6, Papuan-7, Papuan-8, Papuan-9, Pathan-1, Pathan-2, Piapoco-1, Piapoco-2, Pima-1, Pima-2, Polish-1, Punjabi-1, Punjabi-2, Punjabi-3, Punjabi-4, Quechua-1, Quechua-2, Quechua-3, Relli-1, Relli-2, Russian-1, Russian-2, Saami-1, Saami-2, Samaritan-1, Sardinian-1, Sardinian-2, Sardinian-3, She-1, She-2, Sindhi-1, Sindhi-2, Spanish-1, Spanish-2, Surui-1, Surui-2, Tajik-1, Tajik-2, Thai-1, Thai-2, Tlingit-1, Tlingit-2, Tu-1, Tu-2, Tubalar-1, Tubalar-2, Tujia-1, Tujia-2, Turkish-1, Turkish-2, Tuscan-1, Tuscan-2, Ulchi-1, Ulchi-2, Uygur-1, Uygur-2, Xibo-1, Xibo-2, Yadava-1, Yadava-2, Yakut-1, Yakut-2, Yemenite, Yi-1, Yi-2, Zapotec-1, and Zapotec-2.

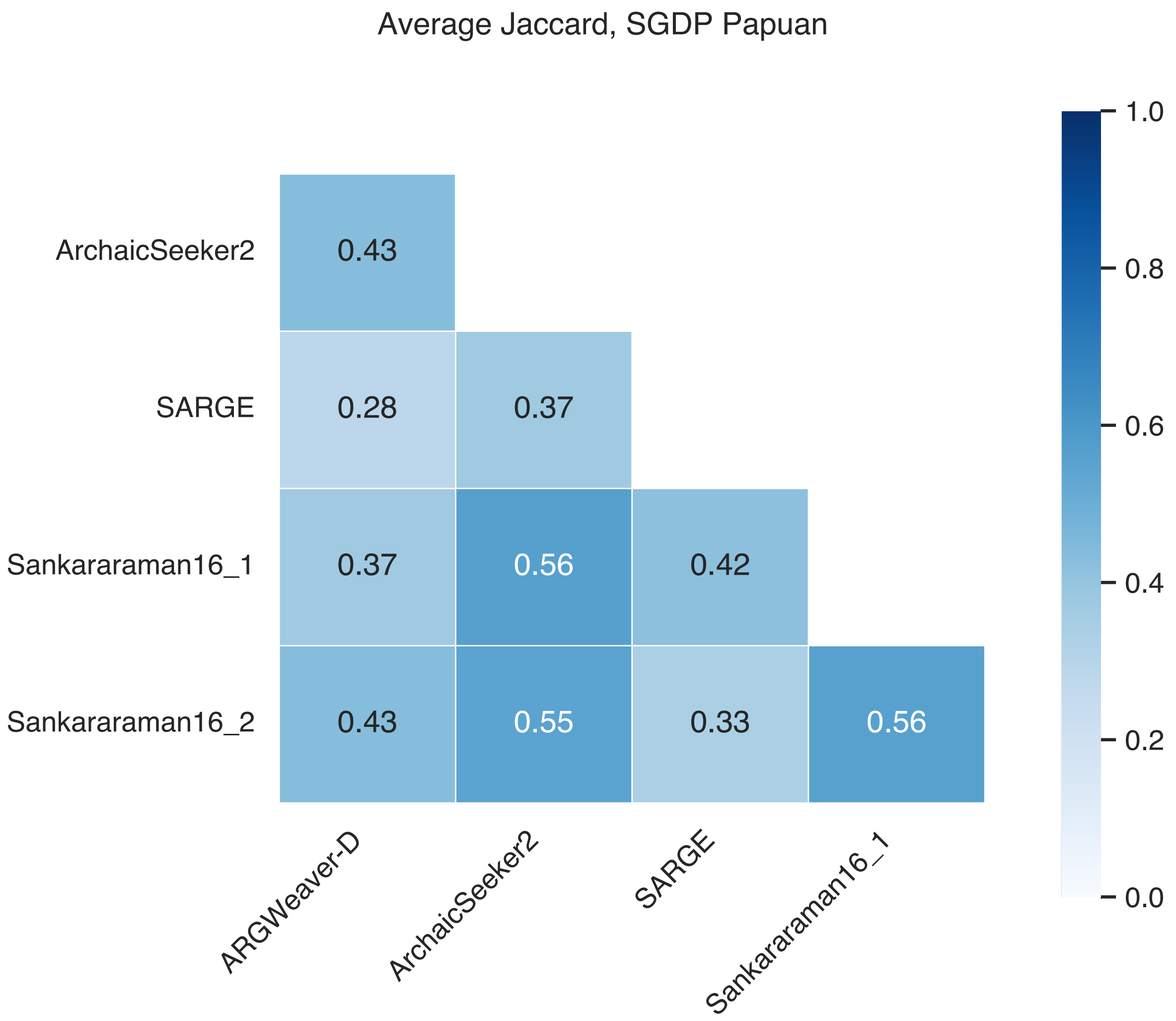


**Supplementary Figure 13:** Raw and normalized Jaccard across introgressed regions in one shared Papuan individual from SGDP (S_Papuan-1). For this comparison, introgression maps included ArchaicSeeker2, SARGE, Sankararaman 2016 (1), and Sankararaman 2016 (2).

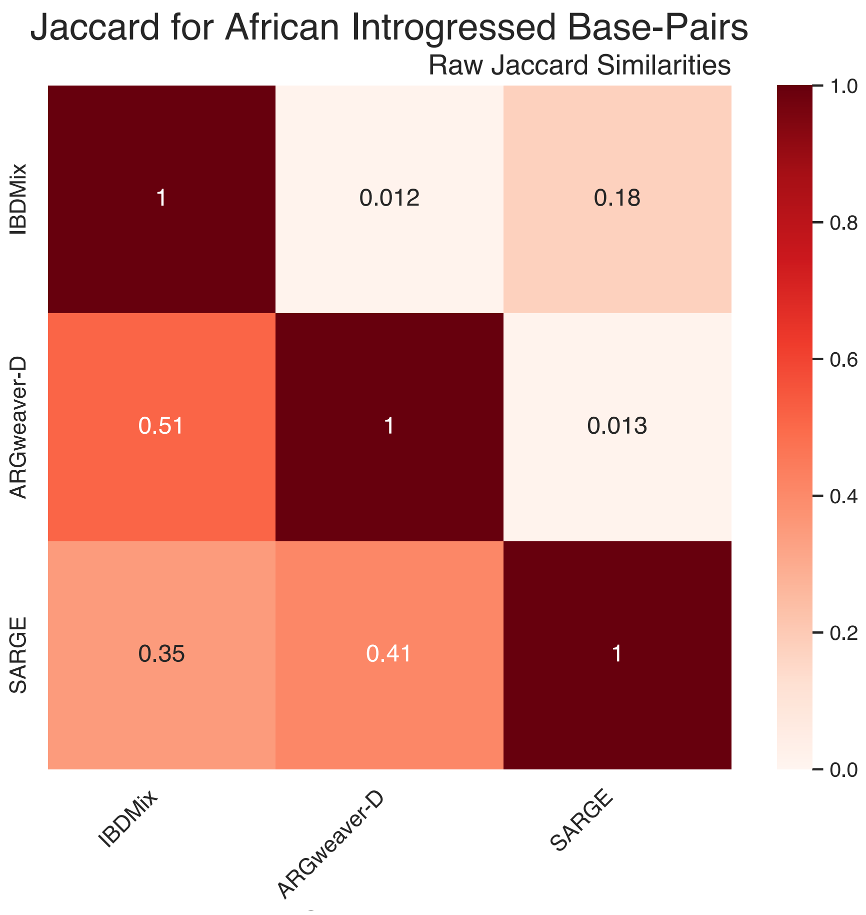


**Supplementary Figure 14: Raw and normalized Jaccard across introgressed regions in African individuals, including admixed individuals in IBDMix.** IBDMix n=661, ARGweaver-D=2, SARGE = 88.

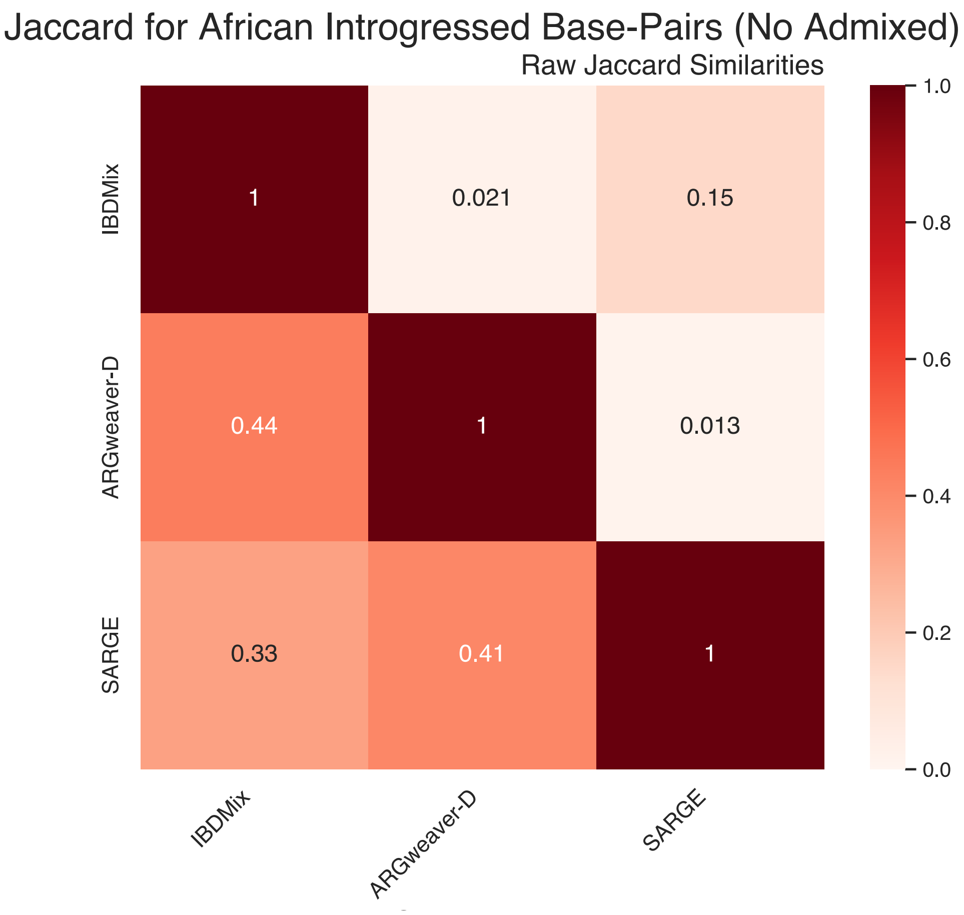


**Supplementary Figure 15: Raw and normalized Jaccard across introgressed regions in African individuals, including admixed individuals in IBDMix.** IBDMix n=504, ARGweaver-D n=2, SARGE n= 88.

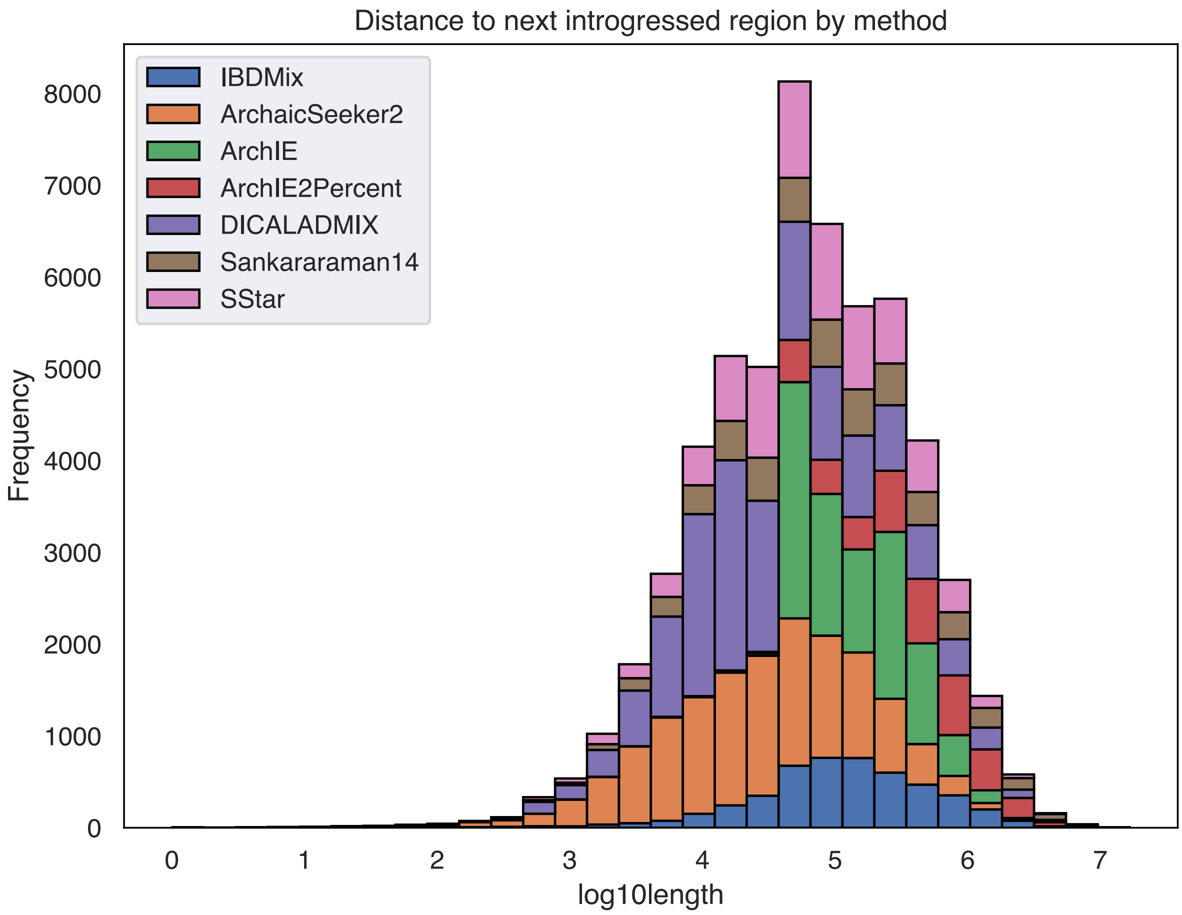


**Supplementary Figure 16:** Distance between introgressed regions across introgression maps with individual-level data.

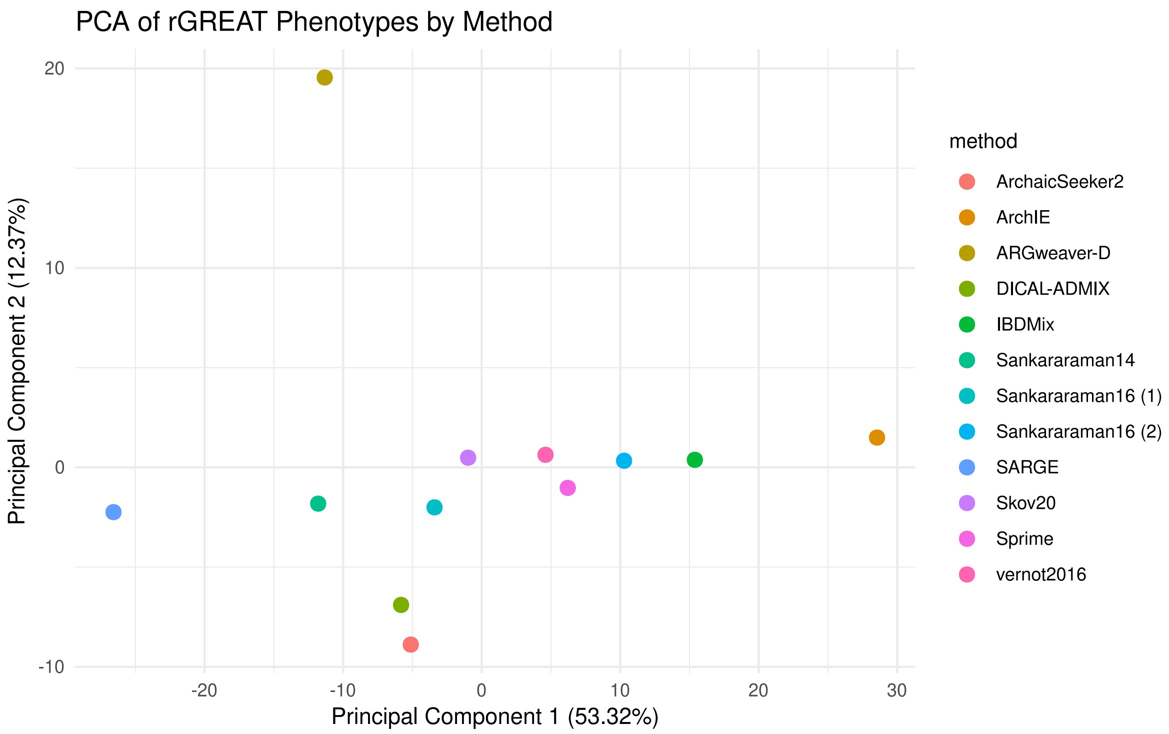

**Supplementary Figure 17: First and second principal components for all method-specific sets of rGREAT phenotypes.** To quantify the similarity between observed enrichments, we computed principal components analysis on the adjusted enrichment p-values for each tested phenotype across all methods. The first principal component accounted for 53% of the explained variance and differentiated all methods, with SARGE and ArchIE as the most distant. Principal component 2 explained 12% of the variance and separated ARGweaver-D, DICAL-ADMIX, and ArchaicSeeker2 from the other methods. ARGweaver-D and SARGE have 116 significant phenotypes shared, the most among sets of shared HPO phenotypes across introgression maps.

| Introgressed loci set | # Phenotypes | # Supported by 3+ separately |
|---|---|---|
| Regions supported by highly supported regions (10+ methods) | 860 | 28 |
| Regions supported by any combination of 10 methods | 871 | 26 |
| Regions supported by all methods, excluding ArchIE and ARGweaver-D | 965 | 25 |
| Regions supported by any combination of 10 methods, all Neanderthal maps as background | 787 | 26 |
| Regions supported by all 12 methods | 356 | 12 |

**Supplementary Table 1:** Number of significant phenotypes associated across varying sets of highly supported introgressed regions

**Method**

| Method | Significant Phenotypes |
|---|---|
| ArchaicSeeker2 | 0 |
| ArchIE | 230 |
| ArgweaverD | 46 |
| IBDMix | 1 |
| Sankararaman14 | 1 |
| Sankararaman16 (1) | 0 |
| Sankararaman16 (2) | 0 |
| SARGE | 1 |
| Skov20 | 4 |
| Sprime | 1 |
| DICAL-ADMIX | 2 |
| Regions supported by any combination of 10 methods | 169 |
| Regions supported by all methods, excluding ArchIE and ARGweaver-D | 48 |
| Regions supported by all 12 methods | 9 |

**Supplementary Table 2:** Number of significant phenotypes associated with each method's introgression map.

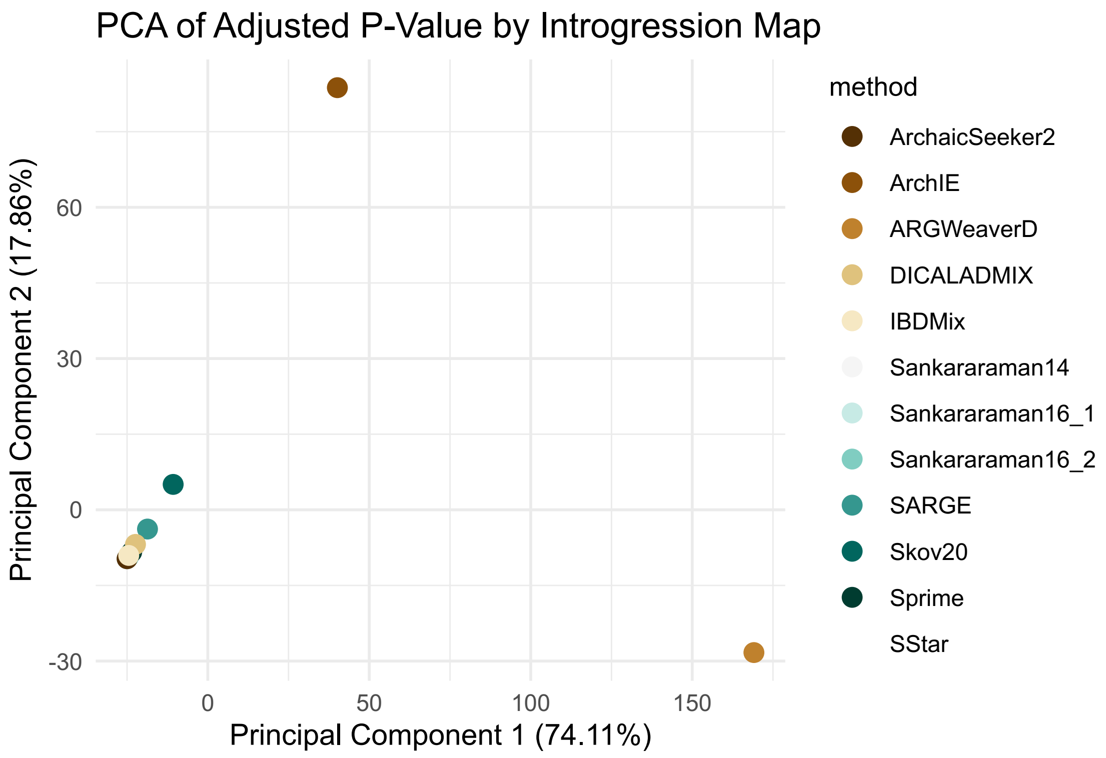


**Supplementary Figure 18**: PCA of adjusted P-values for EnrichR phenotypes associated with each introgression map.

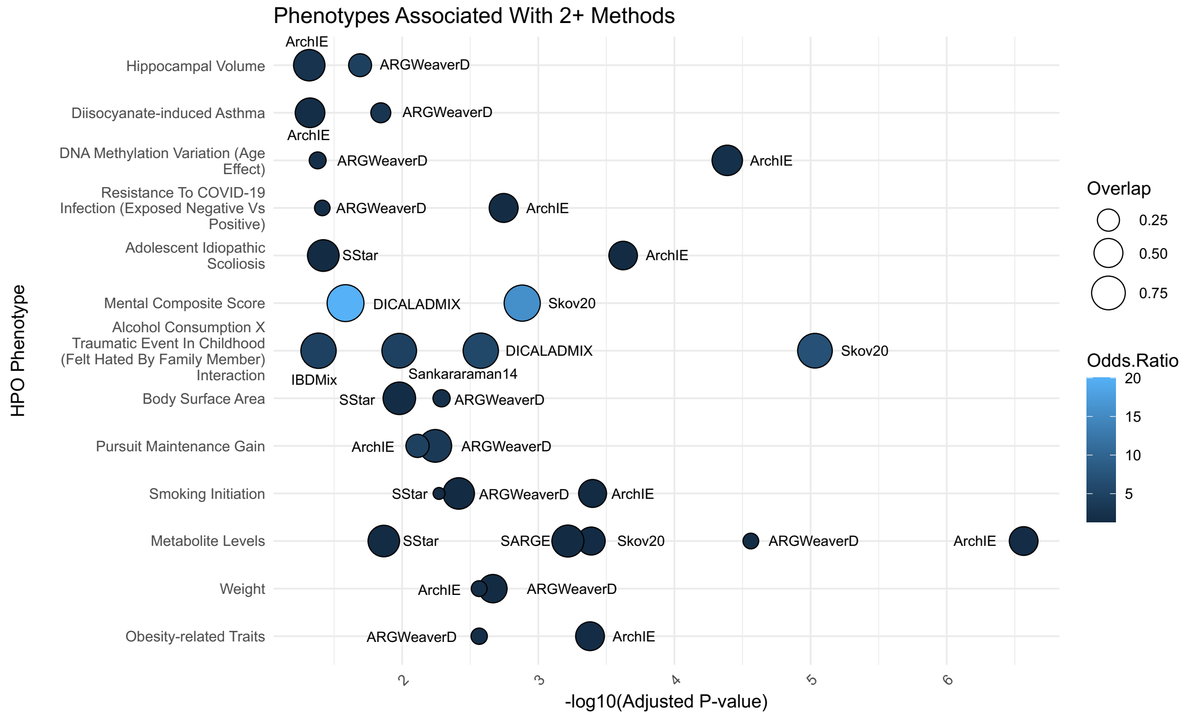


**Supplementary Figure 19**: 13 significant phenotypes associated with multiple introgression maps, with each map as input for EnrichR separately.

| Introgression algorithm | Desert Method | Number and size of desert regions |
|---|---|---|
| S* (Vernot et al, 2016) | Windows across the genome (8-15 Mb, 100kb overlapping steps) in the lower 99th-percentile for average amount of introgression | 6 deserts, 84.4 Mb |
| IBDMix (Chen et al, 2020) | Same as Vernot 2016 | 4 deserts, 64.5 Mb |
| CRF (Sankararaman et al, 2016) | Windows longer than 10 Mb where archaic ancestry proportion <1/1000 | 33 deserts, 483 Mb |
| Two-state HMM (Skov et al, 2020) | 1-Mb windows with no fragments containing Denisovan/Neanderthal variants | 268 deserts, 540 Mb |
| ArchaicSeeker 2.0 (Yuan et al, 2021) | Two-tailed test, 100-kb windows with rare introgression segments | 6 deserts, 79.1 Mb |

**Supplementary Table 3: Desert regions from previous studies with their respective criteria**

**Supplementary File 1:** Significant HPO phenotypes enriched for each introgression map, along with highly supported regions.

**Supplementary File 2:** Genes associated with each region, along with distances to TSS. These files were generated using getRegionGeneAssociations from rGREAT. Associated genes were computed for each introgression map, along with each set of loci included in Supplementary Table 1.
